# High-throughput Automated Muropeptide Analysis (HAMA) Reveals Peptidoglycan Composition of Gut Microbial Cell Walls

**DOI:** 10.1101/2023.04.17.537164

**Authors:** Ya-Chen Hsu, Pin-Rui Su, Lin-Jie Huang, Kum-Yi Cheng, Chun-hsien Chen, Cheng-Chih Hsu

## Abstract

Peptidoglycan (PGN), a net-like polymer constituted by muropeptides, provides protection for microorganisms and has been one of the major targets for antibiotics for decades. Researchers have explored host-microbiome interactions through PGN recognition systems and discovered key muropeptides modulating host responses. However, most common characterization techniques for muropeptides are labor-intensive and require manual analysis of mass spectra due to the complex cross-linked PGN structures. Each species has unique moiety modifications and inter-/intra-bridges, which further complicates the structural analysis of PGN. Here, we developed a high-throughput automated muropeptide analysis (HAMA) platform leveraging tandem mass spectrometry and *in silico* muropeptide MS/MS fragmentation matching to comprehensively identify muropeptide structures, quantify their abundance, and infer PGN cross-linking types. We demonstrated the effectiveness of the HAMA platform using well-characterized PGNs from *E. coli* and *S. aureus* and further applied it to common gut bacteria including species of *Bifidobacterium, Bacteroides, Lactobacillus, Enterococcus,* and *Akkermansia.* We thoroughly explored their PGN structures accurately identified muropeptide mono-/multi-mers, and even unambiguously discriminated the structural isomers via the HAMA platform. Furthermore, we found that the cell stiffness may be correlated to the compactness of the PGN structures through the length of interpeptide bridges or the site of transpeptidation within *Bifidobacterium* species. In summary, the HAMA framework exhibits an automated, intuitive, and accurate analysis of PGN compositions, which may serve as a potential tool to investigate the post-synthetic modifications of saccharides, the variation in interpeptide bridges, and the types of cross-linking within bacterial PGNs.

## INTRODUCTION

Enveloping the cytoplasmic membrane, peptidoglycan (PGN) is an essential and distinct component of bacterial cell walls that maintains the defined bacterial shape, preserves cell integrity by counteracting the internal osmotic pressure,^1^ and acts as a scaffold for other cell wall-anchored components such as proteins and teichoic acids.^2^ PGN also provides a protective barrier for microorganisms against environmental threats, contributing to its structural rigidity. PGN is a net-like polymeric structure composed of various muropeptide molecules, with their glycans linearly conjugated and short peptide chains cross-linked through transpeptidation.^3^ The muropeptide (i.e., PGN-repeating unit) comprises *N-*acetylglucosamine (GlcNAc), *N-* acetylmuramic acid (MurNAc) and a 2–5 amino acid peptide stem, where the two saccharides are linked by β-1,4 glycosidic bonds and the peptide chain covalently binds to the lactyl moiety of MurNAc. In Gram-negative bacteria, the peptide stem generally contains L-alanine, D-isoglutamic acid, *meso*-diaminopimelic acid (*m*DAP), D-alanine, and D-alanine, while in Gram-positive bacteria, it contains L-alanine, D-iso-glutamine, L-lysine, D-alanine, and D-alanine. Although the structural composition of PGN is relatively conserved across species, the mature PGN structure remains heterogeneous and species-specific. This structural diversity is attributed to chemical modifications of the sugar backbone (e.g., *O-*acetylation, *N-*deacetylation, and *N-* glycolylation), alternative amino acids of the peptide stem, interpeptide bridge compositions (e.g., various number of glycine chain), and cross-linking types (e.g., 4-3 linkage, 3-3 linkage, and 2-4 linkages).^4, 5^

In recent years, characterizing muropeptides has become increasingly important, not only because PGN is a target for antibiotic drug design, but also because of the various roles muropeptides play as signaling molecules involved in microbial interaction, antimicrobial release, and host innate immunity.^6–9^ The gut microbiota, consisting of thousands of bacteria, is a key source of PGNs in the host organisms. The signaling functions of gut microbiota-derived PGN fragments in host-microbiota interactions have been extensively studied in inflammation, metabolism, autoimmune diseases, and brain development.^10–13^ Therefore, analyzing the PGN structural composition and its chemical modifications is critical to understand the PGN recognition processes and subsequent activation mechanisms in the immune system.^14^

Nowadays, analytical methods for PGN architecture include HPLC, LC-MS, solid-state NMR, and AFM imaging, enabling a comprehensive study of PGN structure from chemical composition to 3-D architecture.^5^ Online ultra-performance liquid chromatography coupled with electrospray ionization–tandem mass spectrometry (UPLC-ESI-MS/MS) is the primary strategy for muropeptide structural identification, requiring minimal sample amounts and time.^15–17^ However, peak annotation and MS/MS spectra alignment still mostly relies on manual inspection of experienced biological chemists. Previous work published by Bern *et al.* demonstrated on the analysis of muropeptide monomers with MS/MS fragmentation spectra by commercial Byonic peptide identification software.^18^ Later, PGFinder showed consistent and reproducible data analysis of Gram-negative bacterial PGN with MS1 library searching.^19^ Although they both demonstrated great success in identifying muropeptide monomers, the accurate identification of muropeptide multimers and other various bacterial PGN structures still remains unresolved. This is because deciphering the compositions requires MS/MS fragmentation, but it is still challenging to automatically annotate MS/MS spectra from these complex muropeptide structures.

Here, we developed an in-house, high-throughput automated muropeptide analysis (HAMA) platform that simplifies muropeptide structures to sequence format, making it compatible with the (glyco-)proteomics-like “bottom-up” approach. We then established a comprehensive *in silico* MS/MS fragmentation database for muropeptide identification. Using this platform coupled with high-resolution mass spectrometry, we revealed the PGN compositions of common gut bacteria (some of which are Gram-positive), including species from *Bifidobacterium, Bacteroides, Lactobacillus, Enterococcus,* and *Akkermansia*.

## MATERIAL AND METHODS

### Bacterial strains and cell culture

Bacteria and their respective media used in this work were listed in Supplementary Table 1. Anaerobic bacteria such as *Bifidobacterium*, *Bacteroides, Lactobacillus, Enterococcus,* and *Akkermansia muciiniphila* were cultured anaerobically by using an anaerobic workstation (Whitley DG 250, Don Whitley Scientific Limited, England).

### Peptidoglycan extraction and mutanolysin digestion

The extraction of peptidoglycan was performed using a previously described method, with some modifications.^9, 16^ Ten milliliters of the overnight culture that has reached the stationary phase were harvested and lysed by 2% sodium dodecyl sulfate (SDS) solution in 0.1 M Tris/HCl (pH 7.0). The SDS lysate was boiled for 30 minutes at 100 □ in a heating block and spun down at 10,000 rpm for 5 minutes. In order to wash out SDS thoroughly, the pellet was resuspended with 1 mL of H_2_O, centrifuged at 10,000 rpm for 5 minutes, and removed supernatant. After washed with H_2_O three times, the pellet was resuspended in 750 μL of H_2_O and sonicated for 30 minutes. The suspension was spiked with 750 μL of 0.1 M Tris/HCl solution (pH 7.0) and treated with DNase and RNase at 37 □ in a shaker for 1 hour. After the removal of residual nucleic acids, the suspension was subsequently treated with pronase to digest cell wall-bound proteins (final concentration of 0.1 μg/mL for 16 hours at the same conditions). The pronase-treated cell walls were washed twice with H_2_O before wall teichoic acids were released by 1 mL of 1 N HCl incubation for 4 hours. Insoluble peptidoglycans were washed with H_2_O until pH 5-6 and resuspended in 200-500 μL of 12.5 mM sodium dihydrogen-phosphate buffer (pH 5.5) to an OD_600_ of 3.0. One hundred microliters of insoluble peptidoglycan suspensions were digested overnight with 1 μL of mutanolysin solution (1.000 U/mL in H_2_O). Following mutanolysin inactivation (100 □ boiling for 3 minutes) and centrifugation (5 minutes at 10,000 rpm), the supernatants were spiked with 50 μL of reduction solution (10 mg/mL sodium borohydride in 250 mM borate buffer at pH 9.0). After 20 minutes incubation at room temperature, the reduction reaction was stopped by adding 1 μL of phosphoric acid (98%), resulting in pH 2-3. Reduced muropeptide samples were subsequently analyzed by UPLC-MS/MS.

### UPLC-MS/MS analysis of muropeptides

UPLC-MS/MS analysis was performed by a Dionex UltiMate 3000 UHPLC system coupled to a Q-Exactive Plus hybrid quadrupole-orbitrap mass spectrometer (Thermo Scientific). Soluble muropeptides were separated by an ACQUITY UPLC CSH C18 column (130 □, 1.7 μm, 2.1 mm X 100 mm), with a solvent A (0.1% formic acid in water) and solvent B (0.1% formic acid in acetonitrile) as the mobile phase. A flow rate of 0.25 mL/min and 1% mobile phase B were applied in column condition. The injection volume of each sample was 5 μL. An elution gradient was run for 25 minutes: starting with 1% solvent B for 0-2 min; 1-20% B (linear), 2-15 min, 20-95% B (linear), 15-17 min; 95% B, 17-19 min; 95-1% B (linear), 19-21 min; 1% B, 21-25 min for re-equilibration. The column temperature was controlled at 52 □ throughout the whole analysis program. A Heat electrospray ionization (HESI) source was operated in positive mode, with parameters automatically optimized under a flow rate of 250 μL/min (a capillary temperature of 300 □, a probe temperature of 300 □, and a spray voltage of 3.5 kV). Data-dependent acquisition (DDA) mode was used in the instrument. A mass range of MS1 acquisition was from 400 to 2,000 *m/z* at a resolution of 70,000 and the top 5 most abundant ions were subjected to higher-energy collision-induced dissociation (HCD) fragmentation with Δ*m/z* 3 isolation window, stepped normalized collision energy (NCE) at 20, 25, and 35 (A.U.), and dynamic exclusion time of 4 sec.

### Data processing and data analysis

LC-MS/MS raw data was converted to an mzXML format using MSConvert (ProteoWizard) and then processed by HAMA for mass deconvolution and PGN identification.

### Mutanolysin digestion assay

Purified peptidoglycan was adjusted to OD_600_ 1.0 in sodium dihydrogen-phosphate buffer and digested with 5 μL of mutanolysin (1.000 U/mL in H_2_O). Absorbance was measured at OD_600_ of time points at 0, 3, 6, 9, and 16 hours in a Synergy H1 microplate reader (BioTek) with constant orbital shaking at 37 °C.

### Immobilization of cultured bacteria for AFM imaging

The method of bacteria immobilization was performed as described previously, with some modifications.^20^ Overnight cultured bacteria were washed twice and diluted to OD_600_ 0.3 with 1x PBS. Freshly cleaved Si wafers were coated by 500 μL of 0.1% (w/v) poly-L-lysine in H_2_O and left overnight at room temperature. Afterward, the substrates were rinsed five times with ultrapure H_2_O and dried with a nitrogen flow. One hundred microliters of the bacteria suspension were dropped on a silicon substrate, left for 10-20 minutes, and then rinsed three times in H_2_O bath. The bacteria were dried onto the substrate with flowing nitrogen and rehydrated again in 1x PBS bath for liquid imaging.

### Atomic force microscopy (AFM) imaging

Cultured bacteria images were acquired in 1x PBS bath at room temperature by AFM (Dimension Icon, Bruker) with ScanAsyst^TM^. Data were collected by the PeakForce Quantitative NanoMechanics (QNM) mode with qp-CONT probes (spring constant: 0.25 N/m, resonant frequency: 30 kHz, NANOSENSORS^TM^). Images were acquired at a scan rate of 25 μm/s, an applied force of 500 pN, and with a resolution of 256 × 256 pixels per image frame. For mechanical analysis, the approaching part of the curves were fitted with the Derjaguin-Muller-Toporov (DMT) model. DMT modulus of each bacterium was recorded with 16 × 16 curves on top of the bacteria with the area of 250 × 250 nm^2^.

## RESULTS

### HAMA platform: a High-throughput Automated Muropeptide Analysis for Identification of PGN Fragments

Identification and structural analysis of PGNs are time-consuming and challenging due to the complex compositions and modifications. Herein, we developed a high-throughput and automated platform for PGN structural analysis inspired by the (glyco-)proteomic “bottom-up” approach. However, unlike proteins are 1-D amino acid sequences, the PGN is composed by various muropeptides that are intertwined by their glycans and peptide stems. Hence, we first simplified the muropeptide structures as 1-line muropeptide sequences while preserving their complex linkage compositions (Figure 1a, b). Based on the prior knowledge of all possible compositions and modifications of the bacteria species, we used *DBuilder* to build the species-specific database. Subsequently, we analyzed the MS data with *Analyzer* along with the possible *b-* and *y-* ions of CID/HCD fragmentation generated *in silico*. The above-mentioned automated identification processes are compiled in the HAMA platform, a MATLAB-based software with a user-friendly graphic user interface (Supplementary Figure 1 and Additional Files) that includes three parts: *DBuilder*, *Analyzer*, and *Viewer*.

**Figure 1.**
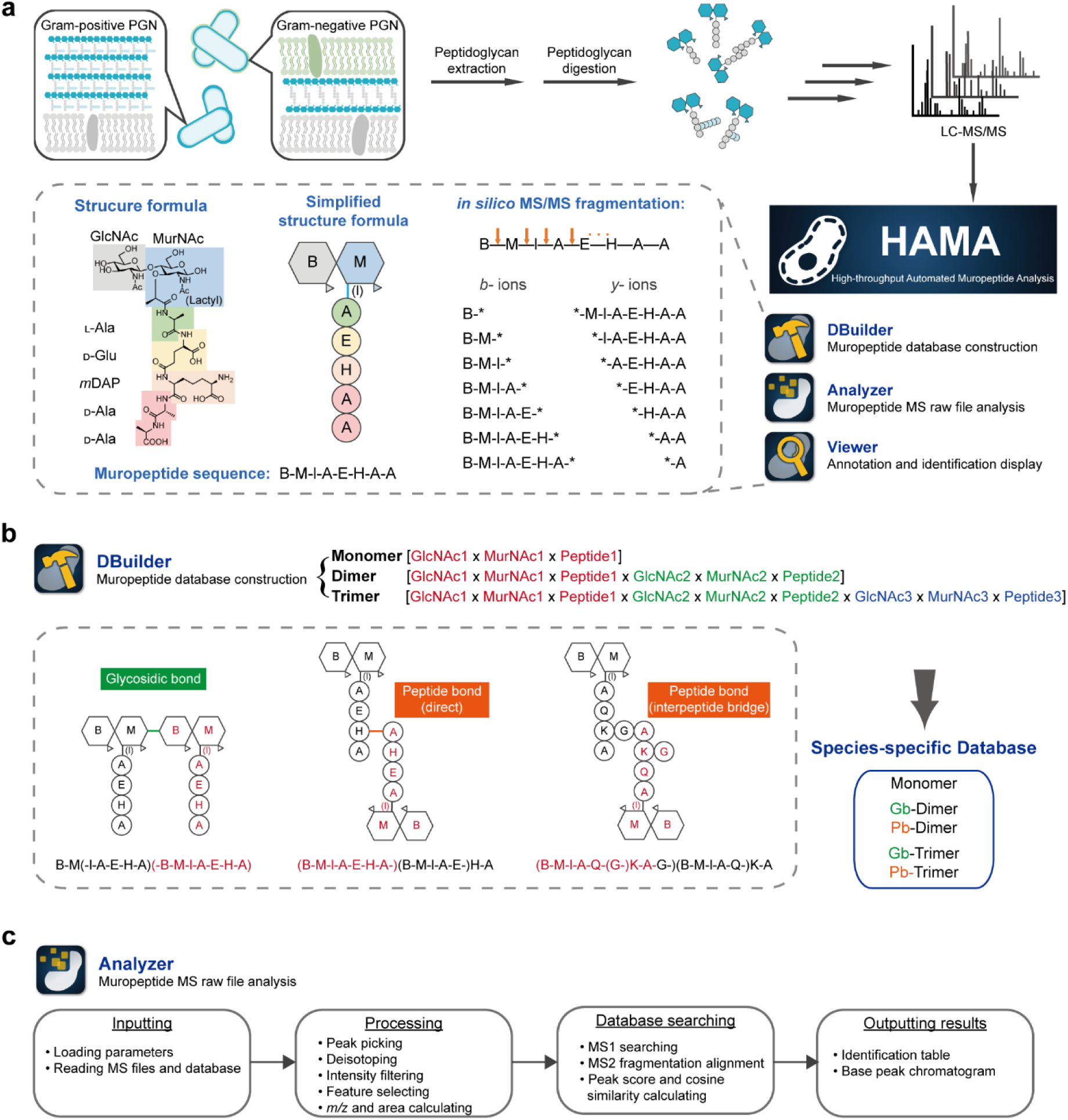
Schematic representation of the high-throughput automated muropeptide analysis (HAMA) framework. (a) The peptidoglycans of bacteria were extracted and purified, followed by mutanolysin digestion. The resulting muropeptide products are analyzed by UHPLC-MS/MS and identified using the HAMA platform. The HAMA strategy involves simplifying muropeptide structures to sequence format, which facilitates the database construction and *in silico* generation of *b-* and *y-* ion fragmentation spectra for matching. Muropeptide symbols: B, *N-*acetylglucosamine; M, *N-*acetylmuraminitol (without lactyl group); l, lactic acid; A, alanine; E, glutamic acid; H, diaminopimelic acid. (b) *DBuilder* constructs a muropeptide database containing monomers, dimers, and trimers with two types of linkage: glycosidic bonds (Gb) and peptide bonds (Pb). For peptide linkages, the direct way is through a direct covalent bond between the penultimate D-Ala of a donor stem and *m*DAP residue in the neighboring acceptor stem, and the indirect way is via an interpeptide bridge branching from the lysine. Donor peptides are labeled in red, and acceptor peptides are labeled in black. (c) The flowchart outlines the LC-MS data processing in *Analyzer*.

(i) ***DBuilder*** *for Species-specific muropeptide database construction.* In order to construct muropeptide databases *in silico*, we named each residue in the muropeptide with a distinct letter and simplified the muropeptide structure into a 1-line sequence format (all representation letters of monosaccharides and amino acids are listed in Supplementary Table 2). Muropeptide sequences were structured with dashes representing peptide/glycosidic bonds and parentheses to discriminate certain subunits. Based on their known (or putative) PGN structures, all possible combinations of GlcNAc, MurNAc and peptide were input into *DBuilder* to generate a comprehensive database that contains monomeric, dimeric, and trimeric muropeptides (Figure 1b). Herein, we constructed PGN multimers using two types of polymerization events: transglycosylation connected via glycosidic bonds, and transpeptidation linked through peptide bonds. In *DBuilder*, the type of peptide linkage used was the common 4-3 cross-link, which could be achieved either directly through a covalent bond between the penultimate D-Ala and the third residue in the acceptor peptide stem, or via an interpeptide bridge. We also considered that the terminal D-Ala (at the fifth position) in the donor stem was not allowed during the transpeptidation reaction. In addition, choosing more possible modifications (anhydro MurNAc residues, deacetyl GlcNAc residues, acetyl MurNAc residues, and amidated iso-Glu) to construct a PGN database could lead to severe mass coincidence issues for MS1 searching in the *Analyzer*. Hence, we set a maximum modification number of six to avoid generating multi-modified muropeptide candidates. Finally, the species-specific muropeptide database was outputted to a *.csv format file, including muropeptide sequences and corresponding chemical formula and theoretical monoisotopic mass.

(ii) ***Analyzer*** *for analysis of muropeptide MS raw files. Analyzer* is an integrated tool for processing MS data and identifying muropeptides. The flowchart is shown in Figure 1c. First, the species-specific database, MS raw file (mzXML format),^21^ and parameters were loaded into *Analyzer*. Parameters were originally set with MS1 range *m/z* 400-2000, retention time range 2- 12 min, mass tolerance of 10 ppm (orbitrap mass spectrometry data), intensity threshold of 1e5, etc., which could be input by users for different experimental conditions. Then, the mzXML file was processed sequentially by peak picking, deisotoping, feature selection, and deconvolution. The observed masses were searched against the loaded database within 10 ppm tolerance. However, in cases where the observed mass matched more than one inferred sequence in MS1 searching, *Analyzer* compared the MS/MS spectra of those MS1 matched features to *in silico* MS/MS fragmentation spectra of corresponding inferred candidates through cosine similarity and matched peak score (MPS, the number of the matched peaks divided by the number of the predicted peaks) calculation. The observed mass was then identified as the sequence with the highest matching score. Lastly, all identified muropeptides were merged into an Excel spreadsheet reporting charge state, molecular weight, retention time, peak intensity, peak area, sequence, main scan number, cosine similarity, score, etc. *Analyze*r also outputted a base peak chromatogram with peak annotations and a result file (in MATLAB data) for *Viewer* input. The entire analysis was completed within a few minutes.

(iii) ***Viewer*** *for annotation and identification display. Viewer*, a visualization tool that allows users to browse extracted ion chromatograms (XICs) of the identified muropeptides and visualize the *in silico* MS/MS matching spectra annotated with *b-* and *y-* ions. In the muropeptide spectra match (PSM) page of each identified muropeptide, *Viewer* lists all inferred candidates and visualizes their individual MS/MS spectral matches to clearly demonstrate the process of scoring in *Analyzer*. Additionally, *Viewer* provides a data sorting function that allows users to classify the identification list by molecular weight, peak area, score (in ascending or descending order), and sequence (in alphabetical order).

### Demonstration of the HAMA Platform Using Well-characterized PGNs of *E. coli* and *S. aureus*

As a proof-of-concept, we demonstrated the HAMA platform using well-characterized peptidoglycans from Gram-negative *Escherichia coli* DH5α and Gram-positive *Staphylococcus aureus* SA113. The typical PGN subunit of *E. coli* is GlcNAc – MurNAc – L-Ala – D-iGlu – *m*DAP – D-Ala – D-Ala, represented as a sequence of B-M-l(-A-E-H-A-A) in the HAMA platform. For *S. aureus*, the classic PGN structure is a disaccharide-pentapeptide with the Gly_5_ interpeptide bridge branching from the lysine, which is simplified as B-M-l(-A-Q-(G-G-G-G-G-)K-A-A. Based on the analyzed input dataset, approximately 70% of the peak area in the base peak chromatograms was assigned to muropeptide signals, which allowed for a comprehensive PGN mapping (Figure 2a, c). The HAMA platform successfully identified *E. coli* and *S. aureus* muropeptides, and their XICs and MS/MS matching spectra could be visualized in *Viewer* (Figure 2b, d). In total, 28 and 89 unique muropeptides were identified in PGNs of *E. coli* and *S. aureus*, respectively. Table 1 lists overviews of the main muropeptide peaks in the chromatograms. Upon examining the entire identification results, we found that some muropeptides were eluted in multiple retention times due to the existence of a few abundant stereoisomers. Additionally, a few monosaccharide muropeptides identified as the loss of GlcNAc could be recognized as in-source fragments by exact co-elution.

**Figure 2.**
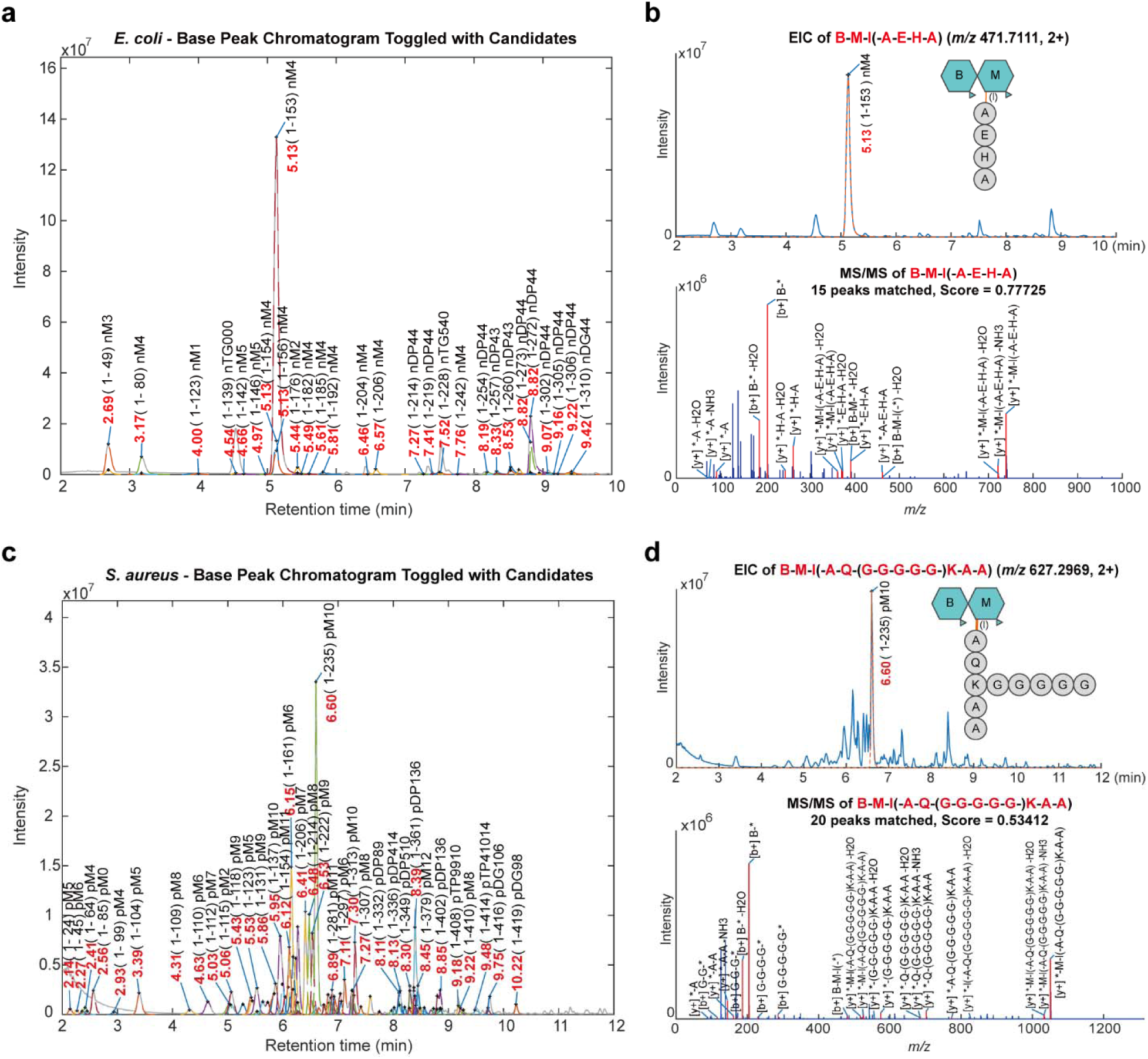
Automated identification of well-characterized peptidoglycans from *E. coli* and *S. aureus* using the HAMA platform. (a, c) Base peak chromatograms showing the muropeptide analysis of *E. coli* and *S. aureus*. The label content includes retention time (in red), feature index, and muropeptide class (in black). (b, d) Extracted ion chromatograms of the most abundant muropeptide and their MS/MS spectra annotated with *b-* and *y-* fragments were visualized in *Viewer*. Muropeptide symbols: B, *N*-acetylglucosamine; M, *N*-acetylmuraminitol (without lactyl group); l, lactic acid; A, alanine; E, glutamic acid; Q, glutamine; K, lysine; G, glycine.

**Table 1.**
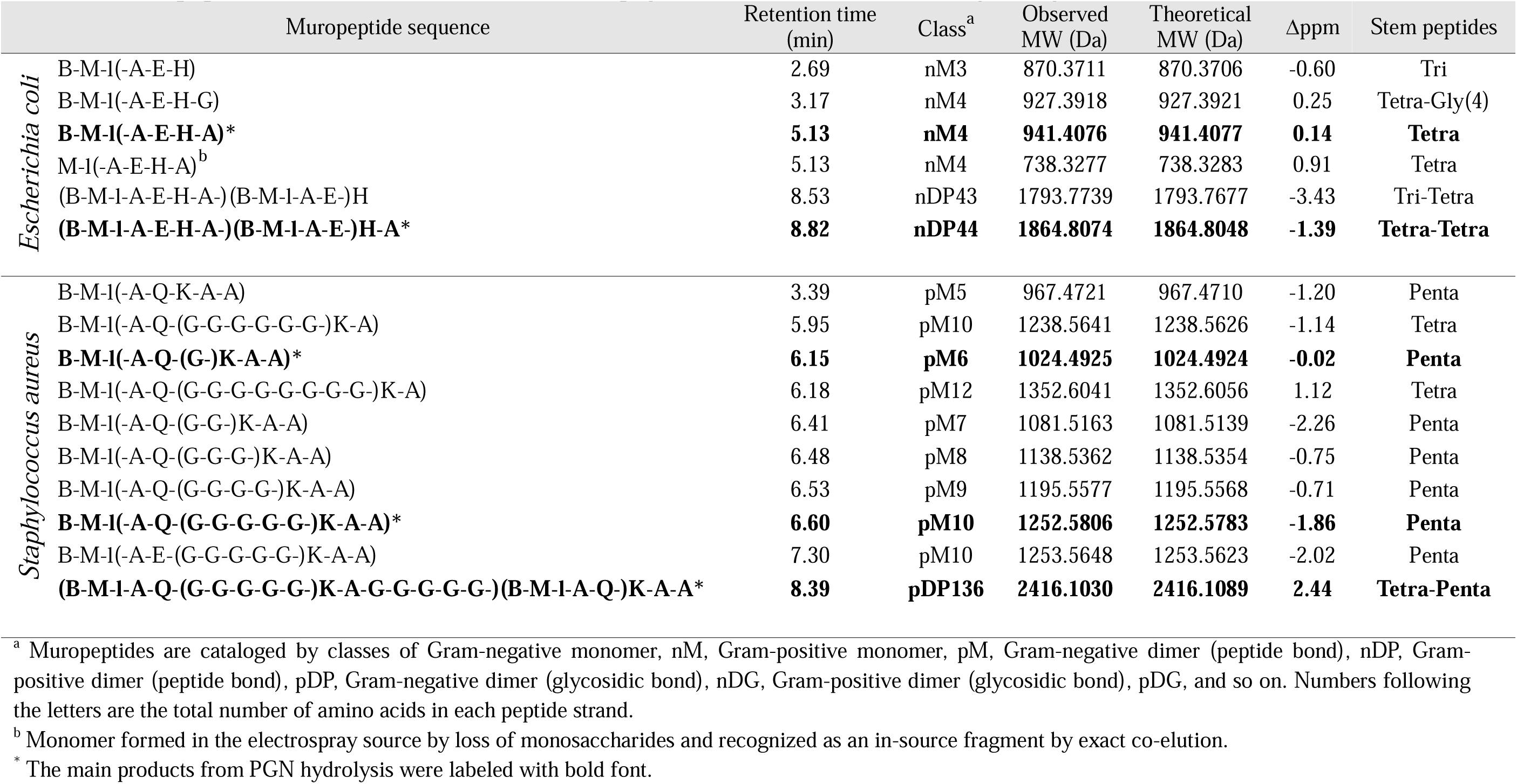
Muropeptides of *Escherichia coli DH5*α / *Staphylococcus aureus* SA113 analyzed by UPLC-MS/MS. **Table 1 - Source data 1** Raw output table of the whole muropeptide identification of *E. coli*. **Table 1 - Source data 2** Raw output table of the whole muropeptide identification of *S. aureus*.

In the *E. coli* data, the most abundant muropeptides were disaccharide-tetrapeptide (Tetra monomer) and disaccharide-tetrapeptide-disaccharide-tetrapeptide (Tetra-Tetra dimer). We also identified two low-abundant muropeptides, B-M-l(-A-E-H-G) and B-M-l(-A-E-H-A-G), in which the fourth and fifth Ala are individually substituted with a Gly. This unique composition has been reported in strain *E. coli* Nissle 1917 from the previous analysis,^16^ but not for DH5α strain used in our study. In the output of *S. aureus* peptidoglycan, the elution profile was the same as what has been previously known: the most abundant monomers were disaccharide-pentapeptide with a (Gly)_5_ bridge and disaccharide-pentapeptide with a Gly bridge. The most abundant cross-linked dimer was disaccharide-tetrapeptide-disaccharide-pentapeptide, which contains a total of ten Gly residues.^16^ We also characterized the known modifications and structural variations within *S. aureus* peptidoglycan, such as *O-*acetylation of MurNAc, the presence of D-iGlu (non-amidated) in the stem peptides, and the length variation of the interpeptide bridge. The high-throughput analysis allowed for the identification of monomeric muropeptides consisting of one to nine Gly residues and dimeric muropeptides containing a total of five to fourteen Gly residues in a single analysis. However, due to the insufficient structural information in the peptide backbone provided by the HCD fragmentation spectra, the exact number of Gly residues harbored in each PGN unit could not be determined. Hence, further careful analysis and manual verification are required to confirm the identity of identified muropeptides, particularly for low-score dimers and trimers. Collectively, these results provide valuable insights into the PGN compositions and architectures.

### Characterizing Gut Bacterial PGN Compositions and Resolving Isomeric Muropeptides

Over the past two decades, an increasing number of gut microbial species have been found to be associated with human health. In addition, emerging evidence has supported the notion that gut microbial muropeptides work as signaling molecules that mediate host−microbiome interactions in metabolism, gut homeostasis, and immunity.^22^ However, little has been discussed about the structures of PGN fragments due to the diversity and complexity of gut bacterial PGNs and the lack of an efficient analytical tool.^14^ To date, most of the structural information on gut bacterial peptidoglycan comes from reports published between 1970 to 2000,^23–25^ whereas that of the health-promoting gut microbes discovered in the last two decades have been rarely reported. Therefore, we utilized the HAMA platform to investigate the specific PGN structures of several common gut bacterial species from *Bifidobacterium, Bacteroides, Lactobacillus, Enterococcus,* and *Akkermansia*. We collected LC-MS/MS data of purified peptidoglycans from ten gut bacterial species and identified those with the self-built species-specific muropeptide databases. Overall, the base peak chromatograms of each species showed approximately 70% of peaks were annotated as muropeptides, and these elution profiles reflected the actual composition and organization within the peptidoglycan architecture. Additionally, the structural compositions of these purified peptidoglycans are consistent with previous reports, listed in Table 2. The muropeptide profiles of the species in this study are summarized in Supplementary Table 3-12.

**Table 2.**
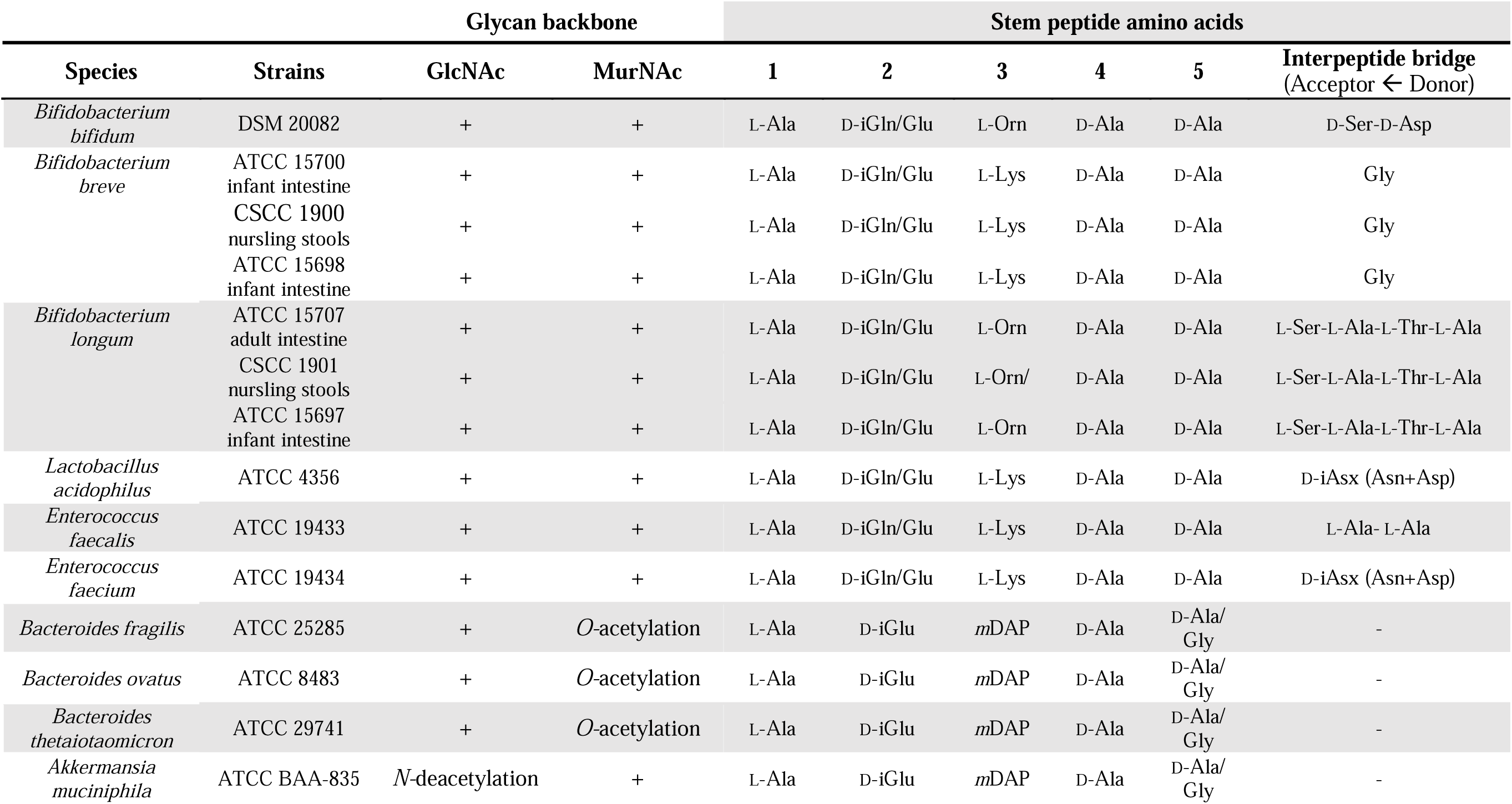
The characterized peptidoglycan types of gut bacteria used in this study.

In the identification results of Gram-positive gut bacterial peptidoglycans, the main muropeptide sequence is GlcNAc – MurNAc – L-Ala – D-iGln – L-Lys – D-Ala – D-Ala, in which the second amino acid of the stem peptide is usually amidated to D-iGln from D-iGlu. This chemical modification is done by the MurT or GatD biosynthetic enzyme and is supposed to control the PGN cross-linking levels, which has recently been demonstrated using labeled PGN stem mimics in certain species.^26^ Another unique feature is L-ornithine (L-Orn) at the third position of the stem peptide. Unlike L-lysine, which is a usual feature of Gram-positive peptidoglycans, L-Orn is a non-proteinogenic amino acid and has been found in the peptidoglycan of certain *Bifidobacterium* species before.^23^ As anticipated, we identified the L-Orn-harbored muropeptides in *B. bifidum* and *B. longum* peptidoglycans. The most species-specific variation among Bifidobacterial PGN was the architecture of the interpeptide bridge, such as Gly, Ser-Asp, and Ser-Ala-Thr-Ala, which corresponded to *B. breve, B. bifidum,* and *B. longum* species. This structural diversity arouses our interest in the relationship between the length of an interpeptide bridge and the physical property of the bacterial cell envelope, which will be discussed in a later section.

In the peptidoglycans of *E. faecium* and *L. acidophilus* (Supplementary Tables 4 and 5), we observed the monomeric structure of GlcNAc – MurNAc – L-Ala – D-iGln – L-Lys – D-Ala – D-Ala, which harbored an interpeptide bridge of asparagine (D-Asn) or aspartate (D-Asp), shortened as B-M-l(-A-Q-(N-)K-A-A) or B-M-l(-A-Q-(D-)K-A-A). In this part, we found that both B-M-l(-A-E-(N-)K) and B-M-l(-A-Q-(D-)K) had identical molecular weights since the mass difference (+ 0.984 Da) between Asn (N) and Asp (D) is the same as between Gln (Q) and Glu (E). This kind of isomeric muropeptides makes identification more complicated, but it can still be addressed by MS/MS *in silico* fragmentation matching under an appropriate separation chromatography. For example, two disaccharide-tripeptides separately eluted at 6.18 min and 7.02 min were found as structural isomers that existed in *E. faecium* peptidoglycan (Figure 3 and Supplementary Table 4). Through *in silico* MS/MS fragmentation matching, these two isomers can be discriminated and identified as the sequences of B-M-l(-A-E-(N-)K) and B-M-l(-A-Q-(D-)K), respectively. To validate the correctness of the automated platform, we also extracted the experimental data of those isomers and manually inspected the MS/MS spectra (Supplementary Figure 2). However, this strategy did not work for the identification of *E. faecalis* peptidoglycan whose interpeptide bridge is composed of two alanine residues (Supplementary Table 3). In this case, two structurally isomeric muropeptides, a disaccharide-tripeptide with a bridge (B-M-l(-A-Q-(A-A-)K)) and a disaccharide-pentapeptide (B-M-l(-A-Q-K-A-A)), have similar *in silico* MS/MS fragmentation patterns, which often leads to misidentification. Nevertheless, the retention time still gave us information to identify them. We annotated the peak at a retention time of 3.66 min as disaccharide-pentapeptide (B-M-l(-A-Q-K-A-A)) by comparing it to an identical sequence that appeared in *L. acidophilus* peptidoglycan with a similar retention time (3.61 min). Therefore, the other peak at a retention time of 6.39 min could be determined as disaccharide-tripeptide with a bridge (B-M-l(-A-Q-(A-A-)K)), which had a longer retention time than the linear structure one.

**Figure 3.**
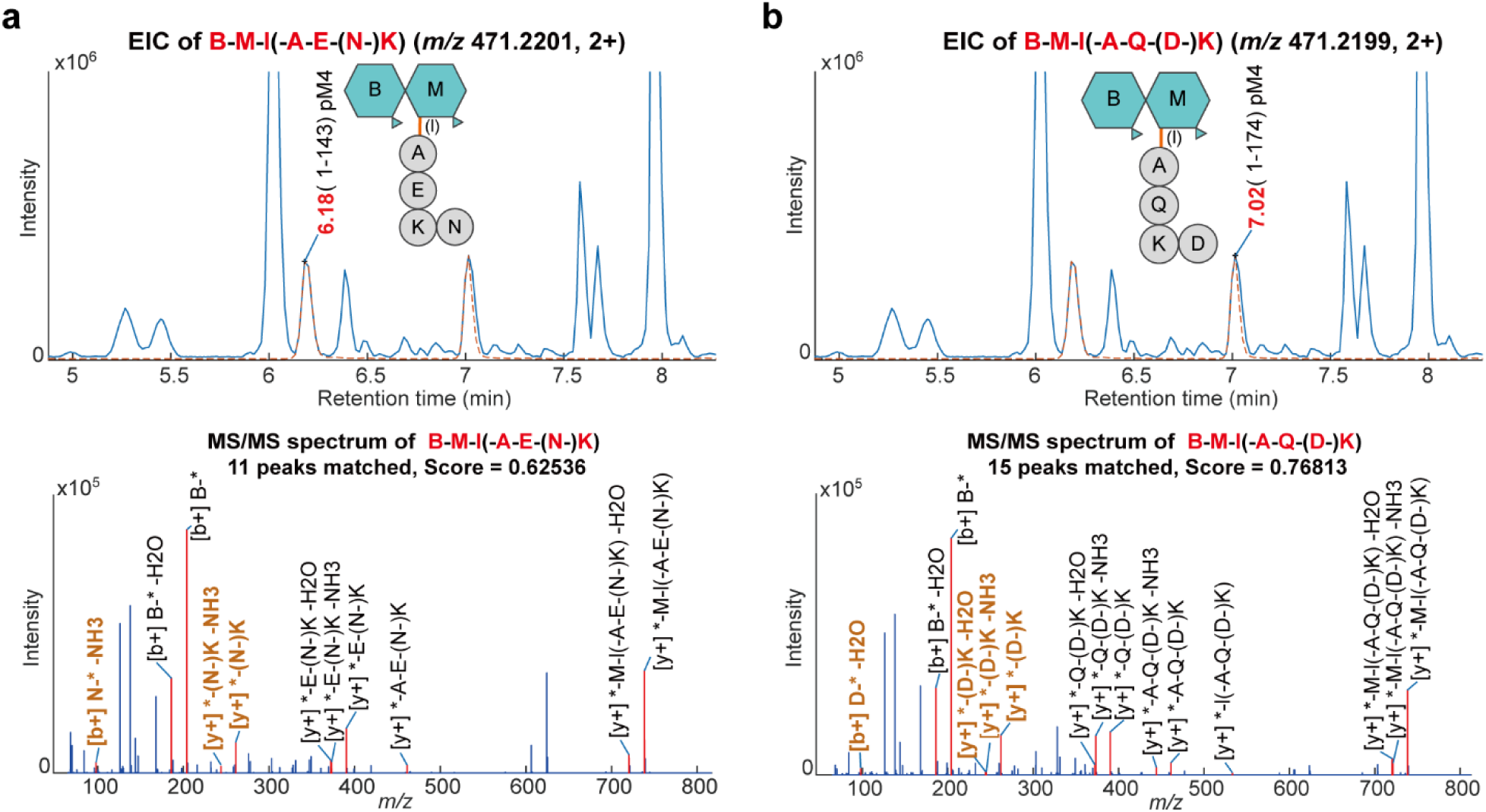
Resolving isomeric muropeptides by *in silico* MS/MS fragmentation matching. Two isomeric muropeptides with the same parent ion, *m/z* 471.22, were identified as two disaccharide-tripeptides: (a) B-M-l(-A-E-(N-)K eluted at 6.18 min, and (b) B-M-l(-A-Q-(D-)K eluted at 7.02 min. The sequence of each isomer was determined using *in silico* MS/MS fragmentation matching, with the identified sequence having the highest matching score. The key MS/MS fragments that discriminate between two isomers are labeled in bold brown. Muropeptide symbols: B, *N*-acetylglucosamine; M, *N*-acetylmuraminitol (without lactyl group); l, lactic acid; A, alanine; E, glutamic acid; Q, glutamine; K, lysine; N, Asparagine; D, Aspartic acid.

Apart from Gram-positive bacteria, we also analyzed the peptidoglycan of several anaerobic Gram-negative bacteria, including *Bacteroides fragilis*, *Bacteroides ovatus*, *Bacteroides thetaiotaomicron*, and *Akkermansia muciniphila* (Table 2 and Supplementary Table 9-12). Structurally, the general stem peptide of Gram-negative peptidoglycan is L-Ala – D-iGlu – *m*DAP – D-Ala – D-Ala, with diaminopimelic acid in the third position being a representative feature. The peptidoglycan structures we identified were consistent with previously published ones, with *O-*acetyl-MurNAc found in *Bacteroides* species and *N-*deacetyl-GlcNAc (GlcN) in *A. muciniphila*.^27, 28^ The output analysis showed that around 56-66% of the total muropeptides in *Bacteroides* species’ peptidoglycans contained *O*-acetylated MurNAc, while approximately 87% of the total muropeptides in *A. muciniphila* peptidoglycan contained de-*N-*acetylated GlcNAc. The high occurrence of *N-*deacetylation in *A. muciniphila* peptidoglycan suggests that *A. muciniphila* might possess a homolog of oxidative stress-induced PGN deacetylase (PgdA) found in *Helicobacter pylori*.^29–31^ Chemical modifications to the disaccharide backbone are known to provide resistance to lysozyme and protect bacteria against enzymatic attack from the host innate immune system.^32, 33^

### Inferring PGN Cross-linking Types Based on Identified PGN Fragments

The substrates and catalyzed enzymes involved in muropeptide cross-linking have been targets for antibiotic development and antimicrobial resistance studies.^34^ Consequently, we sought not only to identify the PGN structures but also to explore the peptide cross-linking types within these gut bacterial PGNs. In general, there are two types of PGN cross-linkage: 4-3 cross-links generated by D,D-transpeptidases (Ddts) and 3-3 cross-links created by L,D-transpeptidases (Ldts) (Figure 4a). Transpeptidation involves two stem peptides which function as acyl donor and acceptor substrates, respectively. As the enzyme names imply, the donor substrates that Ddts and Ldts bind to are terminated as D,D-stereocenters and L,D-stereocenters, which structurally means pentapeptides and tetrapeptides. During D,D-transpeptidation, Ddts recognize D-Ala^4^-D-Ala^5^ of the donor stem (pentapeptide) and remove the terminal D-Ala^5^ residue, forming an intermediate. The intermediate then cross-links the NH_2_ group in the third position of the neighboring acceptor stem, forming a 4-3 cross-link. Following this distinctive rule of PGN biosynthesis, the possible combinations of 4-3 cross-linked dimers are disaccharide-tetrapeptide– disaccharide-pentapeptide (D45) and disaccharide-tetrapeptide–disaccharide-tetrapeptide (D44). During L,D-transpeptidation, Ldts cleave the terminal D-Ala^4^ residue of the donor stem (tetrapeptide) and generate a peptide bond between the third residue and the NH_2_ group in the third position of acceptor stem.^35^ The possible structures of 3-3 cross-linked dimers are disaccharide-tripeptide–disaccharide-pentapeptide (D35), disaccharide-tripeptide–disaccharide-tetrapeptide (D34), and disaccharide-tripeptide–disaccharide-tripeptide (D33). Hence, we could infer the possible PGN cross-linkage types and the involved enzymes based on the main PGN fragments.

**Figure 4.**
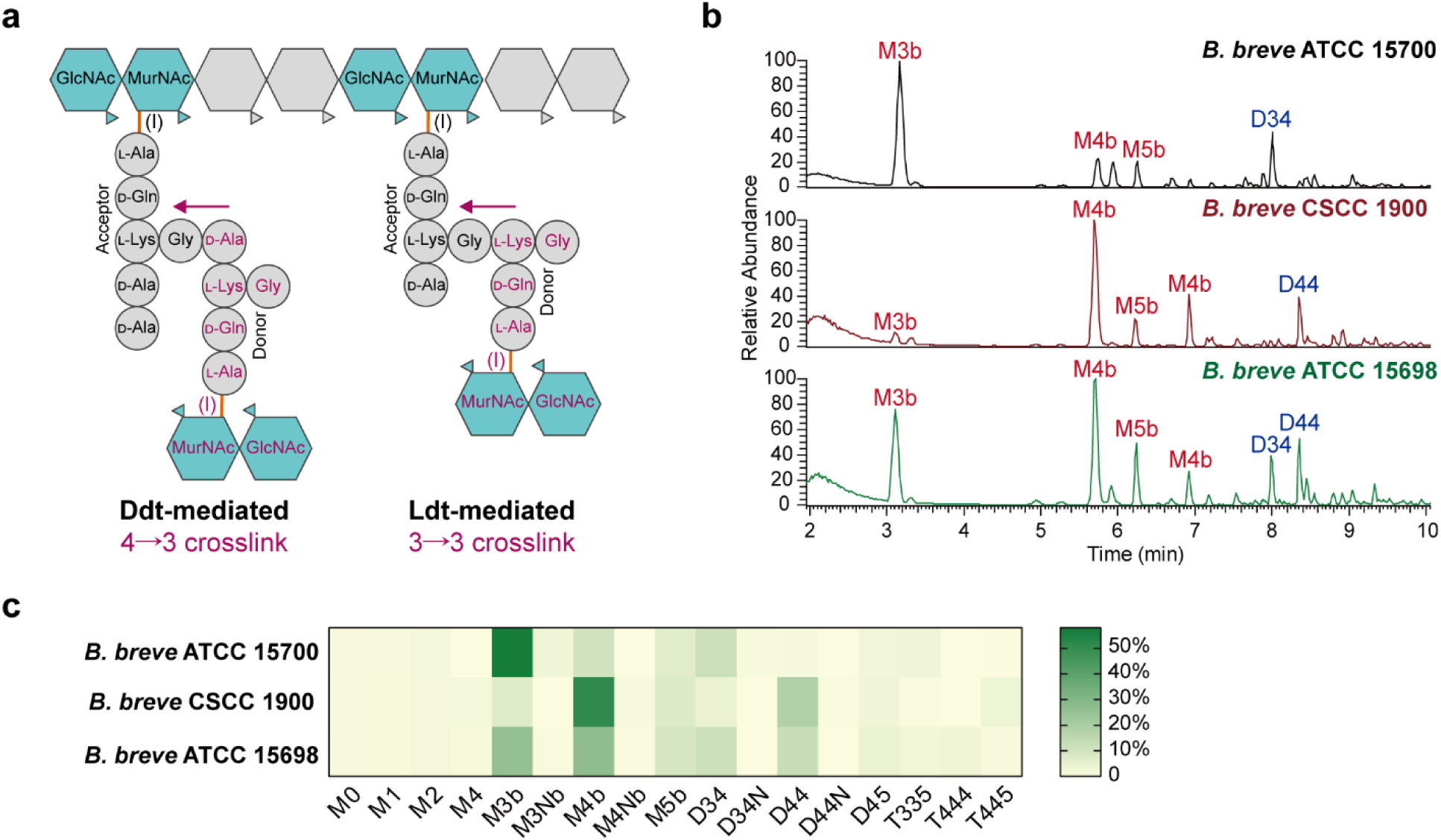
Muropeptide composition analysis of *Bifidobacterium breve* strains. (a) Schematic representation of the two possible cross-linking types in the PGN of *B. breve*: Ddt-mediated 4-3 cross-link and Ldt-mediated 3-3 cross-link. Donor peptide stems are labeled in red. The arrow indicates the direction of cross-links catalyzed by transpeptidases. (b) Base peak chromatograms of muropeptide analysis of *B. breve* ATCC 15700, CSCC 1900, and ATCC 15698 strains. The main peaks were annotated with muropeptide symbols. (c) Heatmap showing the muropeptide compositions (% of total) of the PGN of three *B. breve* strains. Symbols: M, monomer; D, dimer; T, trimer (numbers following the letters indicate the number of amino acids in stem peptides). M0, disaccharide; M1, disaccharide-monopeptide; M2, disaccharide-dipeptide; M4, disaccharide-tetrapeptide; M3b, disaccharide-tripeptide with an interpeptide bridge; M3Nb, disaccharide-tripeptide with an anhydro-MurNAc and an interpeptide bridge; M4b, disaccharide-tetrapeptide with an interpeptide bridge; M4Nb, disaccharide-tetrapeptide with an anhydro-MurNAc and an interpeptide bridge; M5b, disaccharide-pentapeptide with an interpeptide bridge; D34, disaccharide-tripeptide–disaccharide-tetrapeptide with a peptide cross-link; D34N, disaccharide-tripeptide–disaccharide-tetrapeptide with a peptide cross-link and an anhydro-MurNAc; D44, disaccharide-tetrapeptide–disaccharide-tetrapeptide with a peptide cross-link; D44N, disaccharide-tetrapeptide–disaccharide-tetrapeptide with a peptide cross-link and an anhydro-MurNAc; D45, disaccharide-tetrapeptide–disaccharide-pentapeptide with a peptide cross-link; T335, disaccharide-tripeptide–disaccharide-tripeptide-disaccharide-pentapeptide with two peptide cross-links; T444, disaccharide-tetrapeptide–disaccharide-tetrapeptide-disaccharide-tetratapeptide with two peptide cross-links; T445, disaccharide-tetrapeptide–disaccharide-tetrapeptide-disaccharide-pentapeptide with two peptide cross-links. **Figure 4 Source data 1** Heatmap data of Figure 4c.

Take *B. breve* ATCC 15700, CSCC 1900, and ATCC 15698 as examples. The LC-MS profiles showed noticeable differences in muropeptide compositions among the three strains (Figure 4b). The relative muropeptide compositions of each strain are presented in a heatmap (Figure 4c). Obviously, the main muropeptides of *B. breve* ATCC 15700 are disaccharide-tripeptide with an interpeptide bridge (M3b), disaccharide-tetrapeptide with an interpeptide bridge (M4b), and disaccharide-tripeptide–disaccharide-tetrapeptide (D34), whereas the CSCC 1900 strain showed great abundances in disaccharide-tetrapeptide with an interpeptide bridge (M4b), disaccharide-pentapeptide with an interpeptide bridge (M5b), and disaccharide-tetrapeptide–disaccharide-tetrapeptide (D44). These results suggested that 3-3 cross-links and 4- 3 cross-links might be predominant in the peptidoglycans of ATCC 15700 and CSCC 1900 strains, respectively. In the case of ATCC 15698 strain, its peptidoglycan likely contains both types of cross-links since the abundances of M3b and M4b, as well as D34 and D44, are almost equivalent.

Broad-spectrum β*-*lactams are known to inhibit D,D-transpeptidases, such as penicillin-binding proteins (PBPs). However, L,D-transpeptidases are generally insensitive to β*-*lactams and offer alternative cross-links in the PGNs.^36^ Numerous studies have investigated the role of L,D-transpeptidases in the maintenance and remodeling of mature peptidoglycan in organisms such as *E. faecium, C. difficile, E. coli,* and *M. tuberculosis*.^37–42^ Based on the muropeptide compositional analysis mentioned above, we found high abundances of M3/M3b monomer and D34 dimer in the PGNs of *E. faecalis*, *E. faecium, L. acidophilus, B. breve, B. longum,* and *A. muciniphila*, which may be the PGN products formed by Ldts. While the homologs of Ldts in *L. acidophilus, B. breve, B. longum,* and *A. muciniphila* have been supported by genome sequence evidence,^43–46^ biochemical evidence is needed to confirm the existence of L,D-transpeptidases in those species.

### Exploring the Bridge Length-dependent Cell Envelope Stiffness in *B. longum* and *B. breve*

Peptidoglycan, a protective exoskeleton around cells, provides structural integrity to the cell. The porosity of the PGN scaffold, defined by the degree of cross-linking, influences the transport of larger molecules such as proteins. Therefore, modifications to PGN structure are anticipated to significantly affect bacterial cell mechanics and interactions.^47, 48^ Previous evidence has indicated that a high PGN cross-linking level enhances the stiffness of the cell wall material in Gram-positive bacteria.^49^ The PGN-lattice architecture based on the interpeptide bridge length has been investigated using solid-state NMR in *S. aureus*.^50^ However, the effect of the interpeptide bridge length variants on the porosity and stiffness of bacterial cell envelope has little been discussed. Interestingly, the identification table presented above showed that *Bifidobacterium* peptidoglycans have different architecture of the interpeptide bridges among species: a tetrapeptide bridge (Ser-Ala-Thr-Ala) found in *B. longum* and a monopeptide bridge (Gly) found in *B. breve* (Supplementary Figure 3). We wondered whether the length of interpeptide bridges may be related to the bacterial cell envelope’s mechanical properties and hypothesized that the cross-linking with shorter bridges may form a tighter meshwork in peptidoglycan layers (Figure 5a).

**Figure 5.**
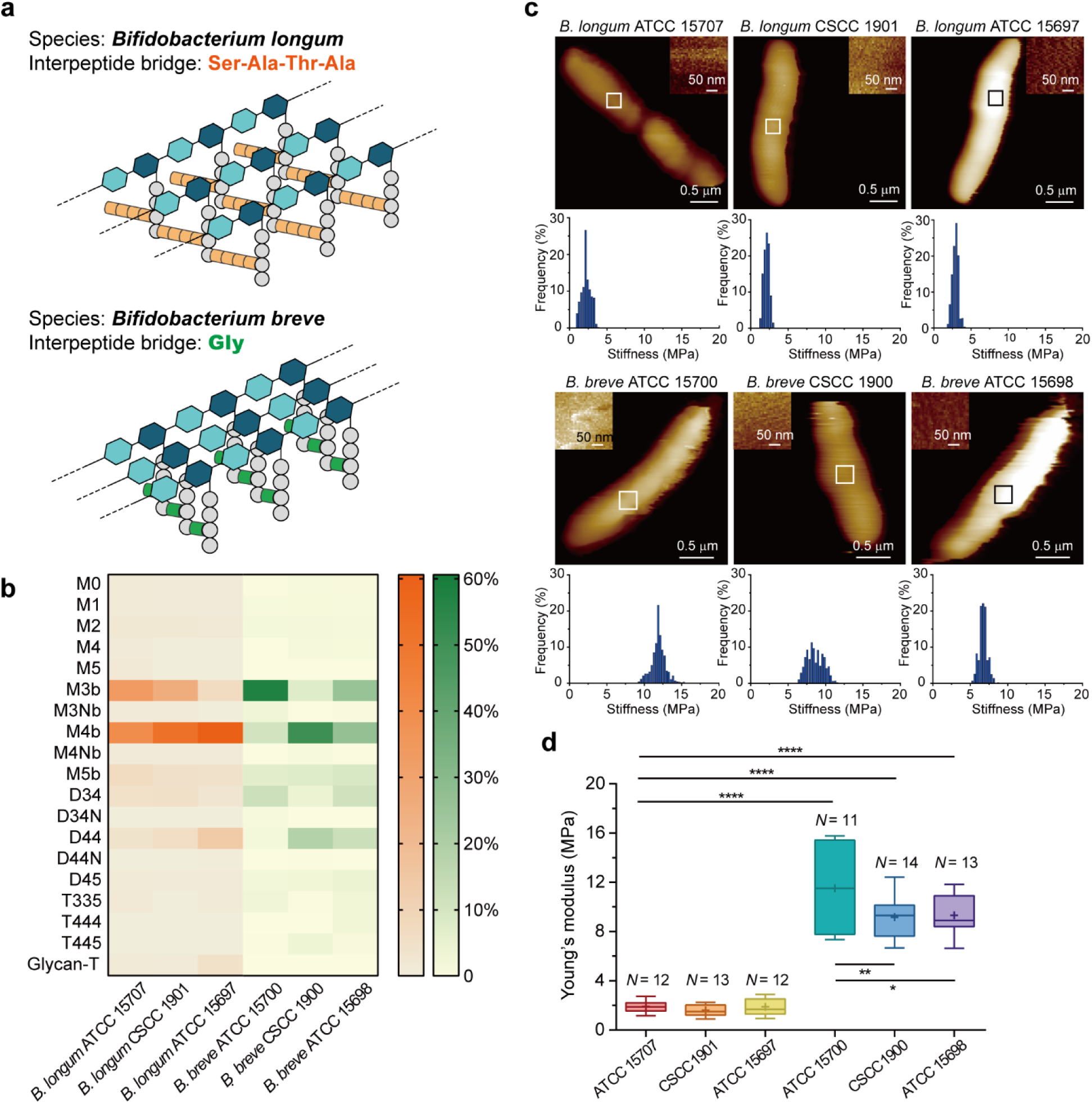
Bridge length-dependent cell envelope stiffness in *B. longum* and *B. breve*. (a) Schematic illustration of the Gram-positive PGN architecture of *B. longum* (orange tetrapeptide bridges) and *B. breve* (green monopeptide bridges). Glycan strands composed of repeating units of the β-1,4-linked disaccharides are cross-linked by interpeptide bridges, forming 3-D peptidoglycan layers. (b) Heatmap displaying the muropeptide compositions (% of total) of the PGNs in *B. longum* and *B. breve* (three strains each). Symbols: M, monomer; D, dimer; T, trimer (numbers indicate amino acids in stem peptides). Description of symbol abbreviations as in Figure legend 4, with the addition of Glycan-T representing trimers linked by glycosidic bonds. (c) AFM imaging of living *Bifidobacterium*. Topographical images in PBS buffer and the inset shows elasticity images from the top of the cell. Distribution of Young’s modulus values corresponding to the elasticity images in the inset. (d) Statistical analysis was performed for each strain, showing the distribution of two groups with the shorter interpeptide bridge corresponded to higher stiffness of cell envelope. Shown here are the mean values (cross), the median, and the 25 and 75% quartiles (boxes) obtained from *N* independent cells over at least three independent experiments. P values were calculated using a one-way ANOVA analysis. *P ≤ 0.05, **P ≤ 0.01, and ****P ≤ 0.0001. **Figure 5 Source data 1** Heatmap data of Figure 5b and AFM data of Figure 5c and 5d.

In order to explore the above relevance, we selected six *Bifidobacterium* bacteria with similar cell morphology as research objects, including three *B. breve* strains and three *B. longum* strains. We summarized the muropeptide compositions of these species (Figure 5b). A slightly higher abundance of peptide-linked dimers in *B. breve* than in *B. longum* implies a higher degree of PGN cross-linking level in *B. breve*. Next, we directly measured the cell stiffness (defined by Young’s modulus, E) of these six bacteria using the single-cell atomic force microscopy (AFM) technique. High-resolution elasticity maps of representative cells were recorded within the central regions to avoid edge effects (Figure 5c). Overall, all *B. breve* were stiffer than *B. longum* with five times higher Young’s modulus values (Figure 5d). Furthermore, *B. breve* also showed a slower enzymatic hydrolysis rate in purified PGNs, implying that the cell wall structure of *B. breve* is characterized by a compact PGN architecture (Supplementary Figure 4). Since *B. breve* strains have shorter interpeptide bridges (one instead of four amino acids in *B. longum*), these data may suggest that the compactness of PGN and the cell stiffness are correlated. Interestingly, among three *B. breve* strains, the ATCC 15700 strain showed significantly stiffer than the other strains, and it also contained more abundant 3-3 cross-linkages in its PGN (i.e., more M3b muropeptides). Computational modeling suggested a more rigid stem peptide in the conformation of L,D-cross-linking, implying the 3-3 cross-linkages may strengthen the PNG layer.^36, 51^ Taken all together, we speculate that a tight peptidoglycan network woven by shorter interpeptide bridges or 3-3 cross-linkages could give bacteria stiffer cell walls. However, it is important to note that cell stiffness is a mechanical property that also depends on PGN thickness, overall architecture, and turgor pressure. These parameters may vary among different bacterial strains. Hence, carefully controlled, genetically engineered strains with similar characteristics will be needed to dissect the role of cross-bridge length in cell envelope stiffness.

## DISCUSSION

The purpose of this study was to develop an automated platform for identifying and analyzing the muropeptide compositions of gut bacterial PGNs. Using high-resolution MS data, we characterized PGN structures of common gut microbes, investigated their PGN cross-linking types, and evaluated the effect of bridge-length variants on the stiffness of bacterial cell walls.

In early 1970s, the amino acid compositions of PGN stem peptides were characterized in many bacteria species, including certain gut microorganisms. At that time, PGNs were purified and hydrolyzed under harsh acidic conditions and then separated by two-dimensional paper chromatography to determine the amino acid sequence. Based on this characterization work, Schleifer and Kandler proposed a bacterial PGN classification system that is still used today.^23^ In the *Bifidobacterium* and *Lactobacillu*s genera, only type A PGNs were identified, representing 4- 3 cross-linking through interpeptide bridges.^25^ However, our study of *B. breve*, *B. longum*, and *L. acidophilus* revealed a significant abundance of M3b and D34 in their muropeptide compositions, implying the incorporation of 3-3 cross-linkage type within PGN network. This cross-linking type is catalyzed by penicillin-insensitive L,D-transpeptidases. We speculate that this finding may be influenced by the comprehensive mass spectrometric approaches we employed or by variations in growth condition. Moreover, we utilized the well-established enzymatic method involving mutanolysin to cleave the β-*N*-acetylmuramyl-(1,4)-*N*-acetylglucosamine linkage, which preserves the original peptide linkage in intact PGN subunits. Importantly, our HAMA platform provides a powerful tool for mapping peptidoglycan architecture, giving structural information on the PGN biosynthesis system. This involves the ability to infer possible PGN cross-linkages based on the type of PGN fragments obtained from hydrolysis. For instance, the identification of 3-3 cross-linkage formed by L,D-transpeptidases (Ldts) is of particular significance. Unlike 4-3 cross-linkages, the 3-3 cross-linkage is resistant to inhibition by β- lactam antibiotics, a class of antibiotics that commonly targets bacterial cell wall synthesis through interference with 4-3 cross-linkages. Therefore, by elucidating the specific cross-linkage types within the peptidoglycan architecture, our approach offers insights into antibiotic resistance mechanisms.

In most Gram-positive bacteria, the interpeptide bridges vary depending on the species and are likely associated with the PGN architectures. Although the chemical structure of PGN units has been extensively studied, the physical structure and 3-D architecture remain open questions.^52, 53^ A previous review by Kim et al. suggested that the length of interpeptide bridges is a key factor in determining different PGN architectures and proposed three PGN-bridge length-dependent architectures based on the orientations of cross-linked peptide stems.^50^ The parallel-stem model, which has the smallest pore size and the highest cross-linking level, is suited for bacteria with long-length bridges. The perpendicular-stem model, with intermediate pore size, is proposed for bacteria with intermediate-length bridges, while the antiparallel-stem model, which has the largest pore size, is for bacteria without bridges. Notably, our study suggested a potential correlation between the cell stiffness and the compactness of bacterial cell walls in *Bifidobacterium* species (Figure 5). *B. longum*, which predominantly harbors tetrapeptide bridges (Ser-Ala-Thr-Ala), exhibits a trend towards lower stiffness, whereas *B. breve*, characterized by PGN cross-linked with monopeptide bridges (Gly), demonstrates a trend towards higher stiffness. These findings suggested that it may be correlated between the increased rigidity and the more compact PGN architecture built by shorter cross-linked bridges. Future studies on the effect of quantitatively changing the PGN-bridge length on stiffness will require intensive research, considering the cross-linking densities, glycan strand lengths, PGN architecture models, and other biomolecules on the cell wall, such as lipoprotein, wall teichoic acids, and lipoteichoic acids.^54–57^ These studies may provide further insight into the physical structure and architecture of PGN, which would contribute to a better understanding of the factors determining bacterial cell wall properties.

In the HAMA platform, simplifying muropeptide structures into sequences has facilitated the generation of *in silico* MS/MS fragments for spectra matching, enabling the structural resolving of isomeric muropeptides. However, there are several limitations to our study design. Our species-specific PGN database is built based on known structures and modifications reported previously. While the identified muropeptides cover approximately 70% of peak area in the base peak chromatograms, certain low-abundant muropeptides composed of uncoded amino acids or saccharides required additional MS/MS identification with manual analysis. For peptide-linked multimers, the current *DBuilder* only builds up 4-3 cross-linked sequences because including the 3-3 cross-link type results in severe misidentifications, as it produces similar *in silico* MS/MS fragmentation patterns as the 4-3 cross-links (about 88-90% similarity). However, we can infer possible PGN cross-linkages based on the type of PGN fragments obtained from hydrolysis.^58^ The primary limitation is that MS/MS spectra generated by HCD fragmentation carry mostly structural information from disaccharide moiety (glyco-oxonium ions) without providing sufficient peptide fragment ions to derive peptide sequence,^59^ making it difficult to accurately identify multimers and determine the position of modifications. Nevertheless, accurate monoisotopic masses acquired from high-resolution mass spectrometry are still informative in peptidoglycan compositional components. Alternative approaches can be considered to elucidate branched structures of muropeptides, such as breaking down PGN with cell wall amidases (cleaving the amide bond between MurNAc and L-Ala residue) to target stem peptide analysis,^60,61^ acquiring MS^3^ spectra to obtain fragmentation maps, or using electron transfer dissociation (ETD) fragmentation mode to leave more peptide fragments.^18^ Additionally, strategies commonly used in glycopeptide identification, such as multi-fragmentation modes (CID, HCD, and ETD), could also be applied to PGN identification to provide more information on peptide sequences and disaccharide structures.^62^ To achieve a more comprehensive identification of muropeptides, future work is needed to generate an expanded database, *in silico-based* fragmentation patterns, and improved MS/MS spectra acquisition.

HAMA is an innovative automated platform that constructs species-specific muropeptide databases, validates structural identification using *in silico* MS/MS analysis, and provides visualized results for efficient and reliable identification of muropeptides, greatly reducing the time-consuming task of manual interpretation of LC-MS/MS data. The HAMA platform has the potential to be a valuable tool for various research fields, including microbiology, pathology, molecular biology, and immunology. It has potential applications in identifying activation ligands for antimicrobial resistance studies, characterizing key motifs recognized by pattern recognition receptors for host-microbiota immuno-interaction research, and mapping peptidoglycan in cell wall architecture studies. With the ease and efficiency offered by HAMA, we believe that muropeptide analysis will become more accessible and contribute to a deeper understanding of cell wall biology.

## ACKNOWLEDGMENTS

This research was supported by Ministry of Science and Technology (MOST), R.O.C. (Grants MOST 108-2636-M-002-008-, 109-2636-M-002-005-, and 110-2636-M-002-014-). Y.-C. H. acknowledged the financial support of “The Program of Research Performance Enhancement via Students Entering Ph.D. Programs Straight from an Undergraduate/Master’s Program” from National Taiwan University. The instrument support from the NTU Mass Spectrometry Platform was acknowledged.

## AUTHOR CONTRIBUTIONS

Y.-C.H., P.-R.S., and C.-C.H. designed the experiments. P.-R.S. and Y.-C.H. built the HAMA software. Y.-C.H. carried out all the MS experiments and analyzed the data. L.-J.H., K.-Y.C., and C.-h.C. designed and carried out the AFM experiments. C.-C.H. supervised the study. Y.-C.H., P.- R.S. and C.-C.H. wrote the paper.

## ADDITIONAL FILES

### Supplementary File

Supplementary file includes a list of bacteria strains, supplementary figures (MS/MS spectra), and supplementary tables (lists of automated identification of gut bacterial muropeptides).

The HAMA software package and *Analyzer* output files are available at: https://drive.google.com/drive/folders/17LVGOm-LEHzNmw7nPULXMEZvg_cnO3k1?usp=share_link

