## Supplementary File for "High-throughput Automated Muropeptide Analysis (HAMA) Reveals Peptidoglycan Composition of Gut Microbial Cell Walls"

### Content


- Supplementary Figure 1. Graphic User Interface (GUI) of HAMA Program.
- Supplementary Figure 2. Tandem MS analysis of two isomeric muropeptides M3b in *E. faecium* PGN.
- Supplementary Figure 3. The dominant lengths of the interpeptide bridges in *B. longum* PGN and *B. breve* PGN.
- Supplementary Figure 4. Peptidoglycan hydrolysis rates and relative muropeptide abundances of different *Bifidobacterium* species.
- Supplementary Table 1. Bacteria strains and cultured media used in this study.
- Supplementary Table 2. Representation letter for each residue of muropeptides.
- Supplementary Table 3. Automated identification of *E. faecalis* ATCC 19433 muropeptides.
- Supplementary Table 4. Automated identification of *E. faecium* ATCC 19434 muropeptides.
- Supplementary Table 5. Automated identification of *L. acidophilus* ATCC 4356 muropeptides.
- Supplementary Table 6. Automated identification of *B. bifidum* DSM 20082 muropeptides.
- Supplementary Table 7-1. Automated identification of *B. breve* ATCC 15700 muropeptides.
- Supplementary Table 7-2. Automated identification of *B. breve* CSCC 1900 muropeptides.
- Supplementary Table 7-3. Automated identification of *B. breve* ATCC 15698 muropeptides.
- Supplementary Table 8-1. Automated identification of *B. longum* ATCC 15707 muropeptides.
- Supplementary Table 8-2. Automated identification of *B. longum* CSCC 1901 muropeptides.
- Supplementary Table 8-3. Automated identification of *B. longum* ATCC 15697 muropeptides.
- Supplementary Table 9. Automated identification of *Bacteroides fragilis* ATCC 25285 muropeptides.
- Supplementary Table 10. Automated identification of *Bacteroides ovatus* ATCC 8483 muropeptides.

- Supplementary Table 11. Automated identification of *Bacteroides thetaiotaomicron* ATCC 29741 muropeptides.
- Supplementary Table 12. Automated identification of *Akkermansia muciniphila* ATCC BAA-835 muropeptides.

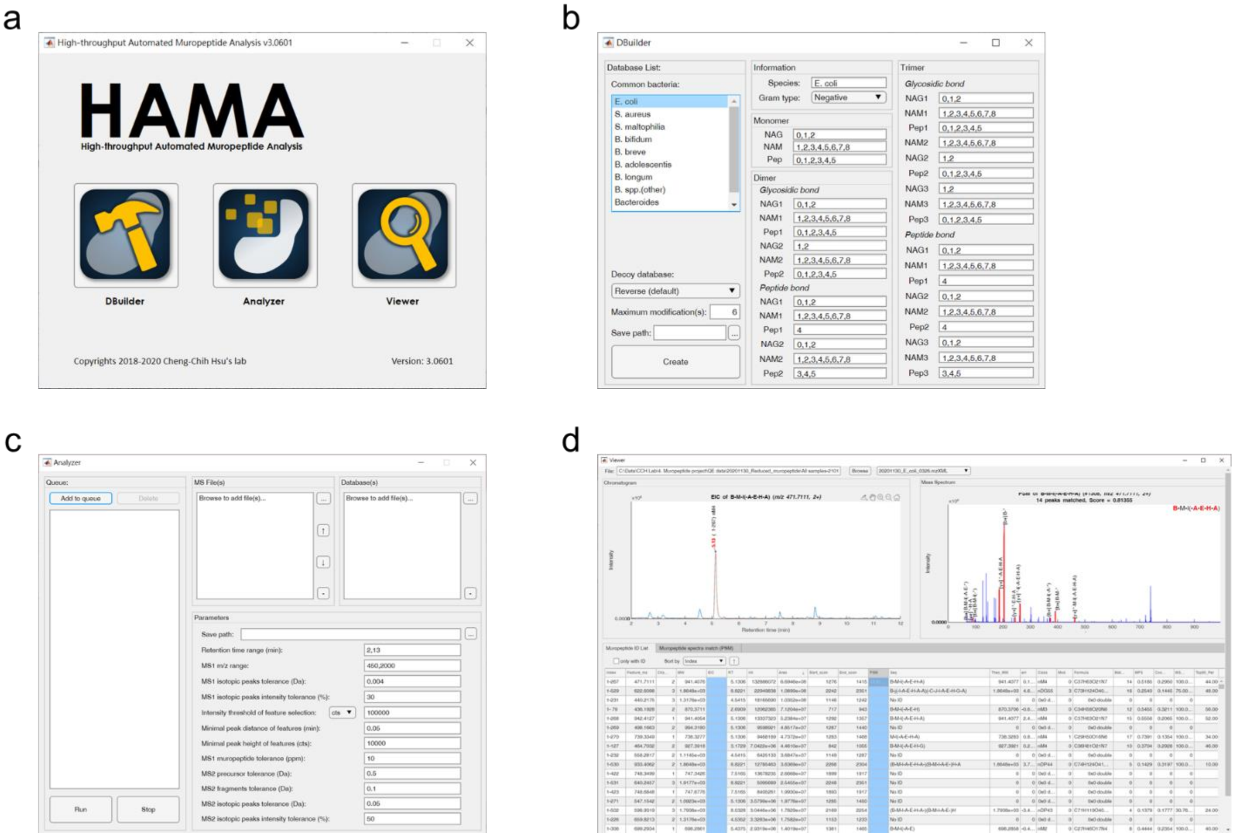


**Supplementary Figure 1. Graphic User Interface (GUI) of HAMA Program.** (a) HAMA provides a user-friendly graphic user interface that makes muropeptide analysis easy to perform. The main GUI comprises three parts: *DBuilder*, *Analyzer*, and *Viewer*. (b) *DBuilder* allows users to construct a species-specific muropeptide database. (c) *Analyzer* offers the flexibility to adjust various parameters for different experimental conditions. (d) *Viewer* enables users to visualize and investigate the identification results.

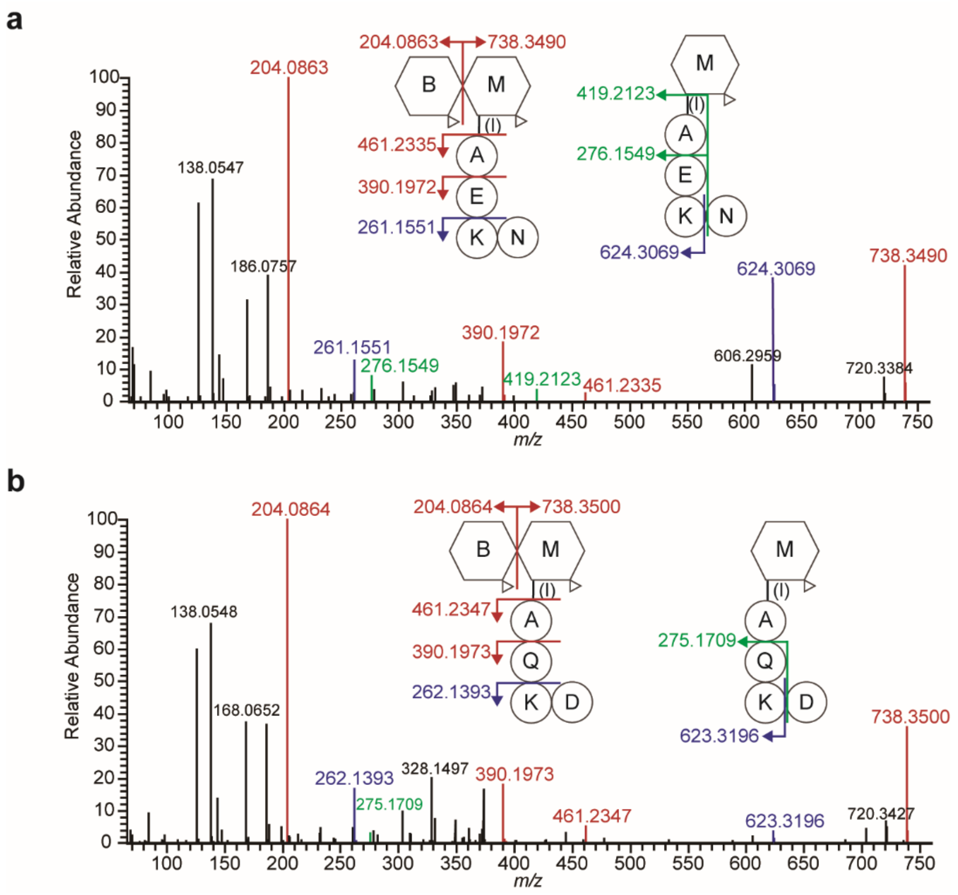


**Supplementary Figure 2. Tandem MS analysis of two isomeric muropeptides M3b in *E. faecium* PGN.** (a) Manually annotated MS/MS spectrum of B-M-l(-A-E-(N-)K (*m/z* 471.2201 at retention time of 6.18 min). (b) Manually annotated MS/MS spectrum of B-M-l(-A-Q-(D-)K (*m/z* 471.2199 at retention time of 7.02 min).

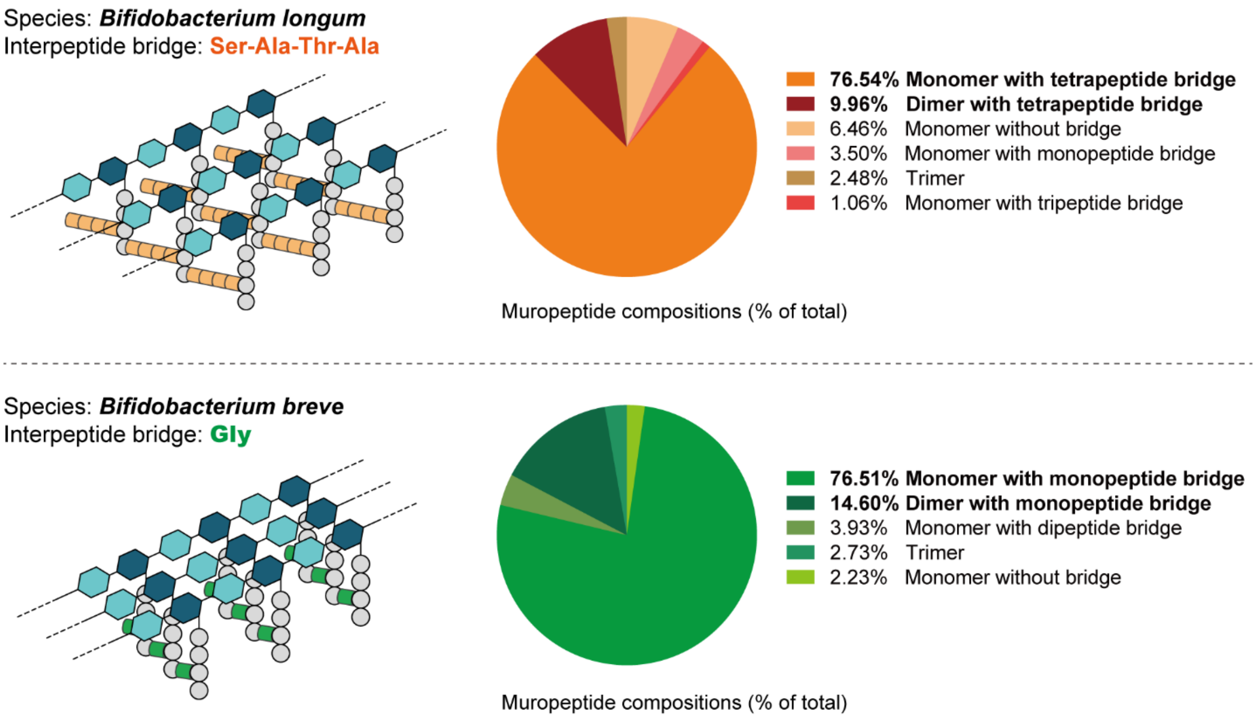


**Supplementary Figure 3. The dominant lengths of the interpeptide bridges in *B. longum* PGN and *B. breve* PGN.** The upper pie chart revealed that approximately 86% of the total muropeptides in *B. longum* PGN were composed of tetrapeptide bridges (Ser-Ala-Thr-Ala). The lower pie chart indicated that approximately 91% of the total muropeptides in *B. breve* PGN were composed of monopeptide bridges (Gly).

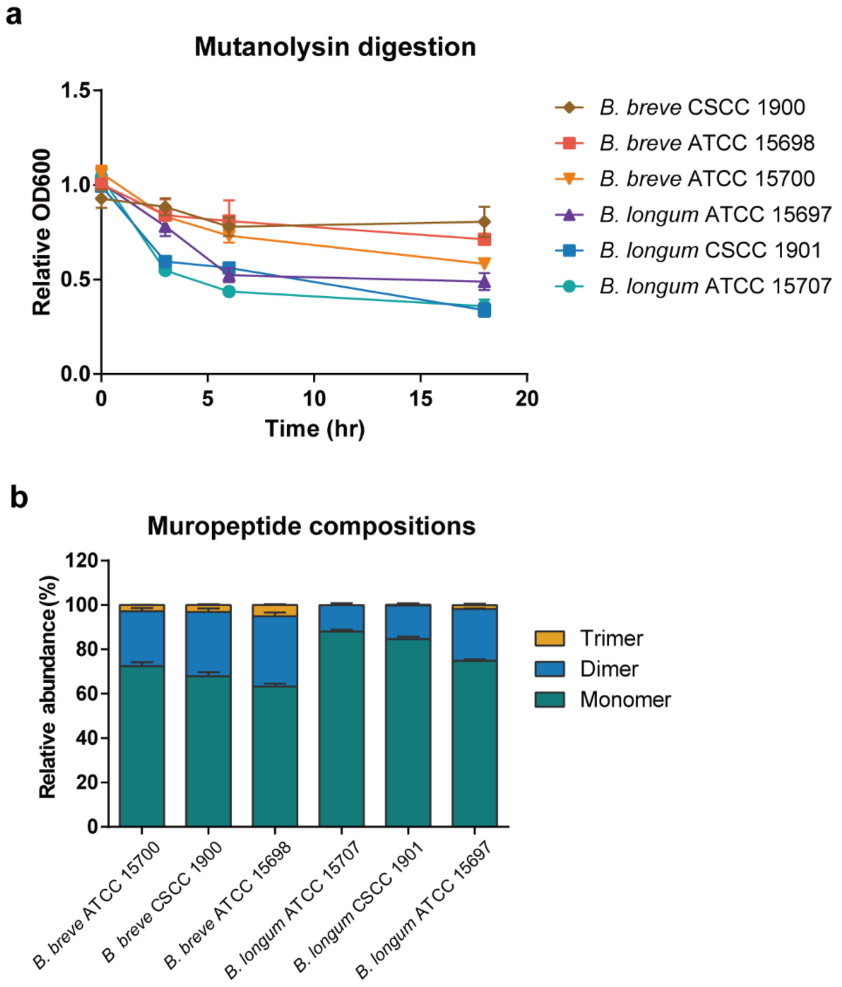


**Supplementary Figure 4. Peptidoglycan hydrolysis rates and relative muropeptide abundances of different *Bifidobacterium* species.** (a) PGN hydrolysis rates of six *Bifidobacterium* strains were measured by monitoring $OD_{600}$ over a 16-hour time course. The results indicate that *B. breve* has a slower PGN hydrolysis rate than *B. longum*. (b) The relative abundance (%) of monomeric, dimeric, and trimeric muropeptides in PGN hydrolysis showed a slightly higher abundance of peptide-linked muropeptides in *B. breve* than in *B. longum*. These experiments were repeated independently at least three times, with similar results.

**Supplementary Table 1. Bacteria strains and cultured media used in this study.**

| | Name of bacterium | Strain | Culture medium |
|---|---|---|---|
| 1 | *Escherichia coli* | DH5$\alpha$ | LB broth |
| 2 | *Staphylococcus aureus* | SA113 | LB broth |
| 3 | *Bifidobacterium bifidum* | DSM 20082 | MRS broth |
| 4 | *Bifidobacterium breve* | ATCC 15700 | MRS broth |
| 5 | *Bifidobacterium breve* | CSCC 1900 | MRS broth |
| 6 | *Bifidobacterium breve* | ATCC 15698 | MRS broth |
| 7 | *Bifidobacterium longum* | ATCC 15707 | MRS broth |
| 8 | *Bifidobacterium longum* | CSCC 1901 | MRS broth |
| 9 | *Bifidobacterium longum* | ATCC 15697 | MRS broth |
| 10 | *Lactobacillus acidophilus* | ATCC 4356 | MRS broth |
| 11 | *Enterococcus faecalis* | ATCC 19433 | RCM broth |
| 12 | *Enterococcus faecium* | ATCC 19434 | RCM broth |
| 13 | *Bacteroides fragilis* | ATCC 25285 | GAM broth |
| 14 | *Bacteroides ovatus* | ATCC 8483 | GAM broth |
| 15 | *Bacteroides thetaiotaomicron* | ATCC 29741 | GAM broth |
| 16 | *Akkermansia muciniphila* | ATCC BAA-835 | GAM broth |

**Supplementary Table 2. Representation letter for each residue in muropeptides.**

| Residue | | 1-Letter code | Chemical Formula | Monoisotopic mass |
|---|---|---|---|---|
| GlcNAc | Native | B | $C_8H_{13}NO_5$ | 203.0794 |
| | Deacetylated | C | $C_6H_{11}NO_4$ | 161.0688 |
| | Acetylated | F | $C_{10}H_{15}NO_6$ | 245.0899 |
| **NAc-muraminitol** (without lactyl group) | Native | **M** | $C_8H_{15}NO_5$ | 205.0950 |
| | Deacetylated | **J** | $C_6H_{13}NO_4$ | 163.0845 |
| | Acetylated | **Z** | $C_{10}H_{17}NO_6$ | 247.1056 |
| MurNAc (without lactyl group) | Native | m | $C_8H_{13}NO_5$ | 203.0794 |
| | Deacetylated | j | $C_6H_{11}NO_4$ | 161.0688 |
| | Acetylated | z | $C_{10}H_{15}NO_6$ | 245.0899 |
| | Anhydrous | U | $C_8H_{11}NO_4$ | 185.0688 |
| | Anhydro/ deacetylated | P | $C_6H_{11}NO_3$ | 145.0739 |
| Lactyl group of MurNAc | | l | $C_3H_4O_2$ | 72.0211 |
| Amino acid | Alanine (Ala) | A | $C_3H_5NO$ | 71.0371 |
| | Asparagine (Asn) | N | $C_4H_6N_2O_2$ | 114.0429 |
| | Aspartic acid (Asp) | D | $C_4H_5NO_3$ | 115.0269 |
| | Glutamic acid (Glu) | E | $C_5H_7NO_3$ | 129.0426 |
| | Glutamine (Gln) | Q | $C_5H_8N_2O_2$ | 128.0586 |
| | Glycine (Gly) | G | $C_2H_3NO$ | 57.0215 |
| | Lysine (Lys) | K | $C_6H_{12}N_2O$ | 128.0950 |
| | Serine (Ser) | S | $C_3H_5NO_2$ | 87.0320 |
| | Threonine (Thr) | T | $C_4H_7NO_2$ | 101.0477 |
| | Diaminopimelic acid (DAP) | H | $C_7H_{12}N_2O_3$ | 172.0848 |
| | Ornithine (Orn) | O | $C_5H_{10}N_2O$ | 114.0793 |

**Supplementary Table 3. Automated identification of *Enterococcus faecalis* ATCC 19433 muropeptides.**

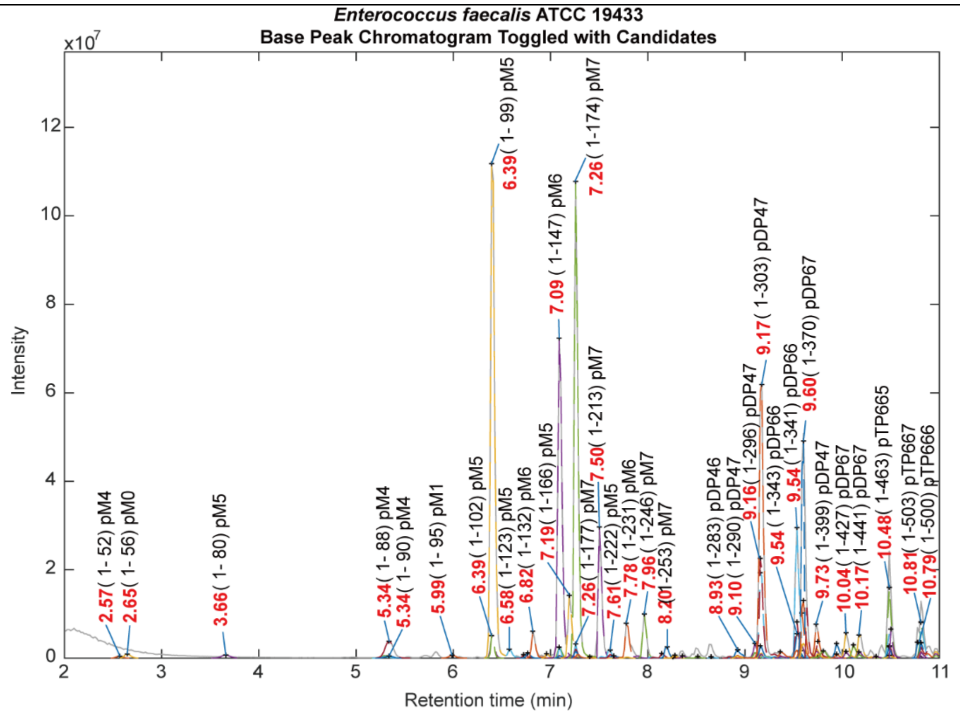


| Muropeptide structure (Main peaks) | RT (min) | Class | Observed MW (Da) | Theoretical MW (Da) | Δppm |
|---|---|---|---|---|---|
| **B-M-l(-A-Q-(A-A-)K** | **6.39** | **Monomer** | **967.4699** | **967.4710** | **1.07** |
| **B-M-l(-A-Q-(A-A-)K-A)** | **7.09** | **Monomer** | **1038.5076** | **1038.5081** | **0.45** |
| B-M-l(-A-E-(A-A-)K | 7.19 | Monomer | 968.4565 | 968.4550 | -1.56 |
| **B-M-l(-A-Q-(A-A-)K-A-A)** | **7.26** | **Monomer** | **1109.5448** | **1109.5452** | **0.40** |
| B-M-l(-A-E-(A-A-)K-A-A) | 7.96 | Monomer | 1110.5330 | 1110.5292 | -3.42 |
| **Tri-Tetra** | **9.17** | **Dimer** | **1987.9670** | **1987.9685** | **0.77** |
| Tetra-Tetra | 9.54 | Dimer | 2059.0038 | 2059.0056 | 0.86 |
| **Tetra-Penta** | **9.60** | **Dimer** | **2130.0409** | **2130.0427** | **0.86** |
| Inferred types of peptide cross-links: 3-3 crosslink /4-3 crosslink | | | | | |

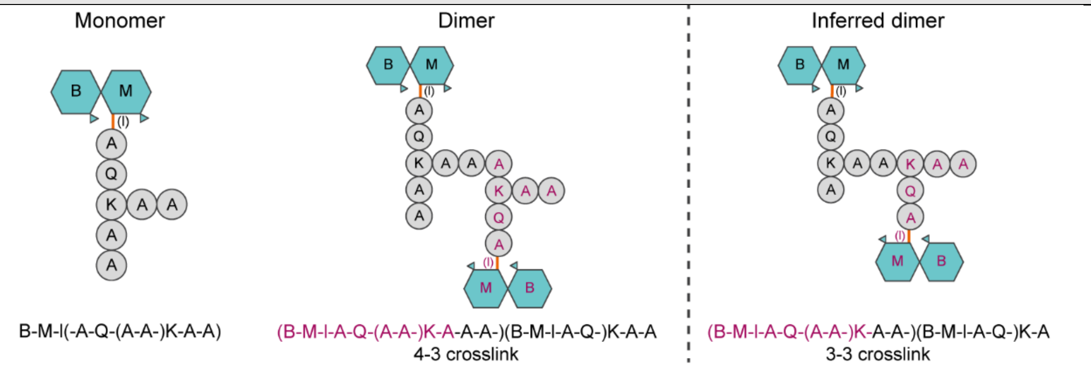


* The main products from PGN hydrolysis were labeled with bold font.

**Supplementary Table 4. Automated identification of *Enterococcus faecium* ATCC 19434 muropeptides.**

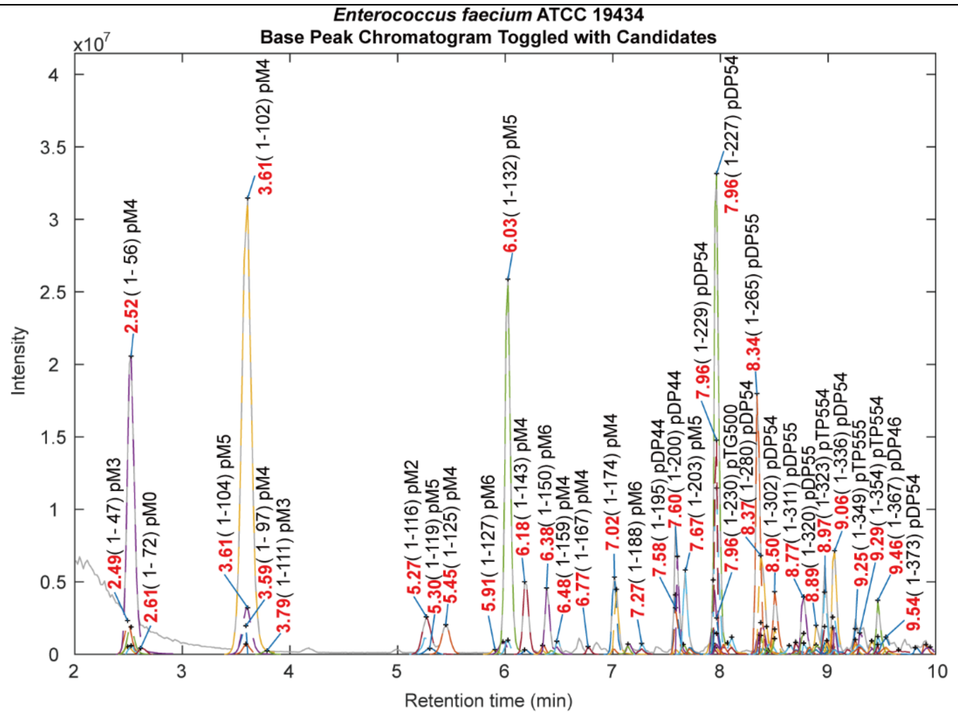


| Muropeptide structure (Main peaks) | RT (min) | Class | Observed MW (Da) | Theoretical MW (Da) | Δppm |
|---|---|---|---|---|---|
| **B-M-l(-A-Q-K-A)** | **2.52** | **Monomer** | **896.4332** | **896.4339** | **0.76** |
| **B-M-l(-A-Q-(N-)K)** | **3.61** | **Monomer** | **939.4397** | **939.4397** | **-0.04** |
| **B-M-l(-A-Q-(N-)K-A)** | **6.03** | **Monomer** | **1010.4765** | **1010.4768** | **0.26** |
| B-M-l(-A-Q-(D-)K) | 7.02 | Monomer | 940.4252 | 940.4237 | -1.62 |
| B-M-l(-A-Q-(D-)K-A) | 7.67 | Monomer | 1011.4622 | 1011.4608 | -1.38 |
| **Tri-Tetra (N, N)** | **7.96** | **Dimer** | **1931.9057** | **1931.9059** | **0.08** |
| **Tetra-Tetra (N, N)** | **8.34** | **Dimer** | **2002.9413** | **2002.9430** | **0.84** |
| Tri-Tetra (D, N) | 9.06 | Dimer | 1932.8914 | 1932.8899 | -0.76 |
| Inferred types of peptide cross-links: 3-3 crosslink /4-3 crosslink | | | | | |

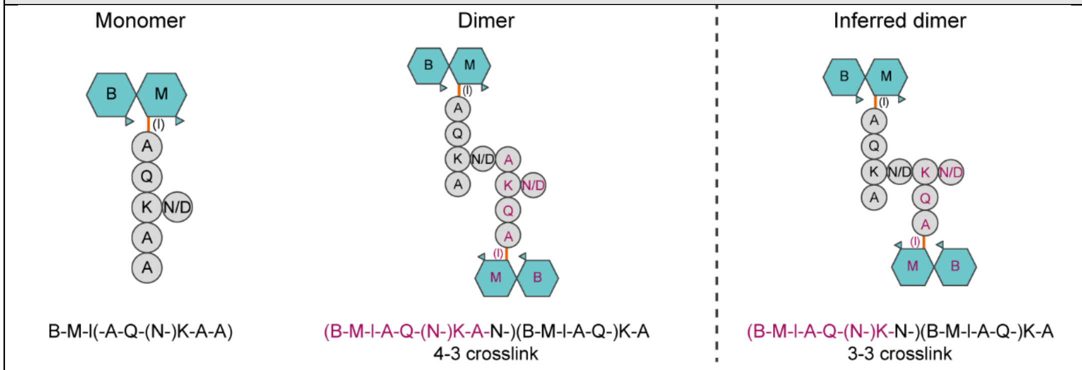


* The main products from PGN hydrolysis were labeled with bold font.

**Supplementary Table 5. Automated identification of *Lactobacillus acidophilus* ATCC 4356 muropeptides.**

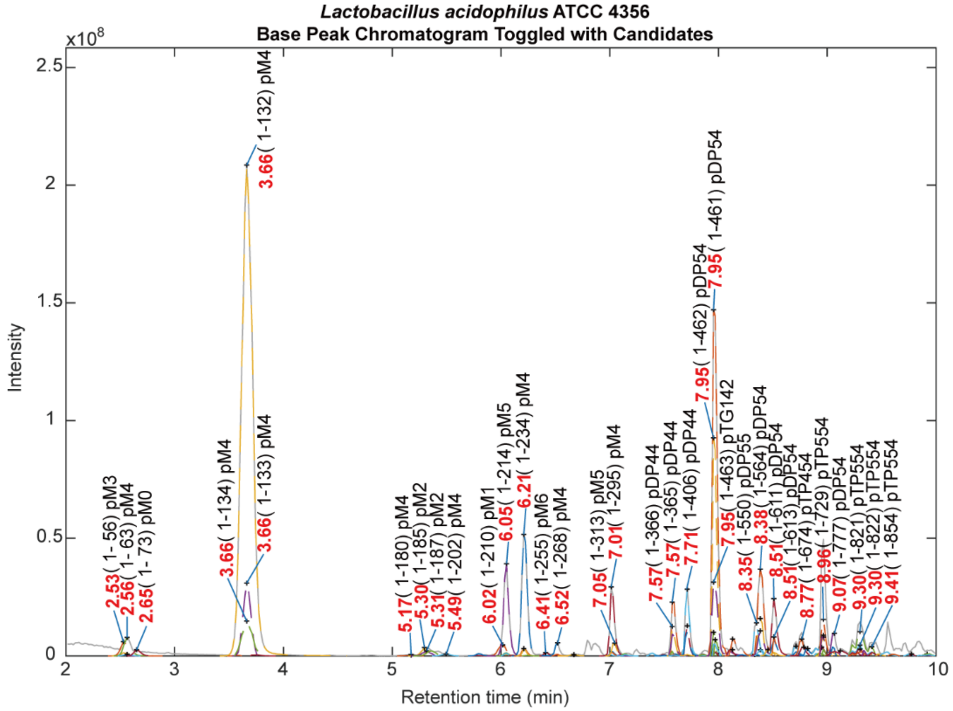


| Muropeptide structure (Main peaks) | RT (min) | Class | Observed MW (Da) | Theoretical MW (Da) | Δppm |
|---|---|---|---|---|---|
| **B-M-l(-A-Q-(N-)K)** | **3.66** | **Monomer** | **939.4368** | **939.4397** | **3.08** |
| B-M-l(-A-Q-(N-)K-A) | 6.05 | Monomer | 1010.4768 | 1010.4768 | -0.05 |
| B-M-l(-A-E-(N-)K) | 6.21 | Monomer | 940.4243 | 940.4237 | -0.62 |
| B-M-l(-A-Q-(D-)K) | 7.01 | Monomer | 940.4245 | 940.4237 | -0.87 |
| Tri-Tetra (N, no interpeptide) | 7.57 | Dimer | 1817.8613 | 1817.8630 | 0.93 |
| **Tri-Tetra (N, N)** | **7.95** | **Dimer** | **1931.8996** | **1931.9059** | **3.26** |
| Tri-Tetra (N, D) | 8.38 | Dimer | 1932.8901 | 1932.8899 | -0.12 |
| Tri-Tetra-Tetra (N, N, D) | 9.30 | Trimer | 2925.3598 | 2925.3561 | -1.27 |
| Inferred types of peptide cross-links: 3-3 crosslink /4-3 crosslink | | | | | |

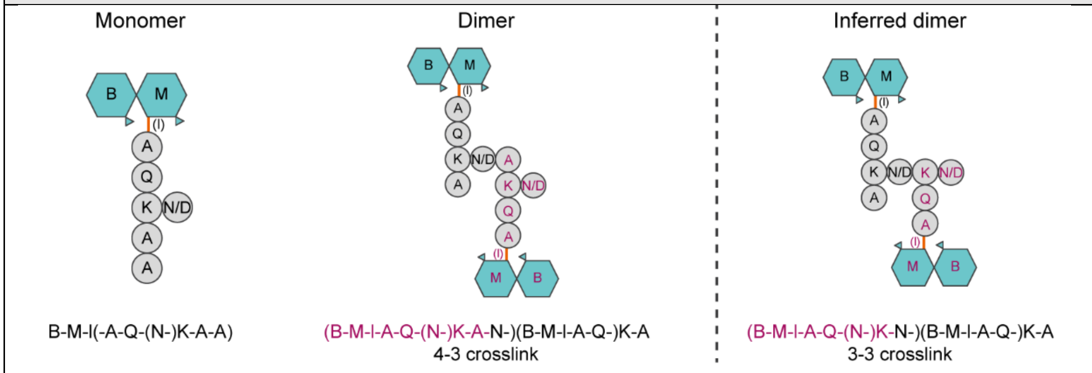


* The main products from PGN hydrolysis were labeled with bold font.

**Supplementary Table 6. Automated identification of *Bifidobacterium bifidum* DSM 20082 muropeptides.**

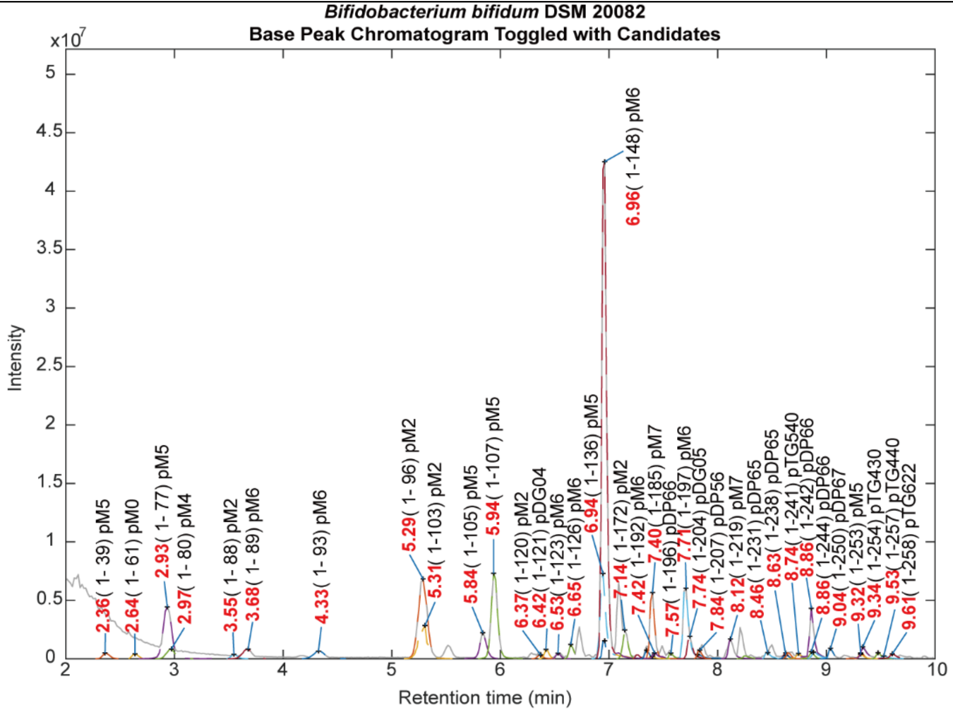


| Muropeptide structure (Main peaks) | RT (min) | Class | Observed MW (Da) | Theoretical MW (Da) | Δppm |
|---|---|---|---|---|---|
| B-M-l(-A-Q) | 5.29 | Monomer | 697.3026 | 697.3018 | -1.15 |
| B-M-l(-A-Q-(D-S-)O) | 5.94 | Monomer | 1013.4430 | 1013.4401 | -2.91 |
| B-M-l(-A-E-(D-S-)O) | 6.94 | Monomer | 1014.4223 | 1014.4241 | 1.78 |
| **B-M-l(-A-Q-(D-S-)O-A)** | **6.96** | **Monomer** | **1084.4775** | **1084.4772** | **-0.28** |
| B-M-l(-A-Q-(D-S-)O-A-A) | 7.40 | Monomer | 1155.5152 | 1155.5143 | -0.78 |
| B-M-l(-A-E-(D-S-)O-A) | 7.71 | Monomer | 1085.4632 | 1085.4612 | -1.85 |
| **Tetra-Tetra** | **8.86** | **Dimer** | **2150.9477** | **2150.9438** | **-1.82** |
| Tetra-Penta | 9.04 | Dimer | 2221.9898 | 2221.9809 | -4.02 |

Inferred types of peptide cross-links: 4-3 crosslink

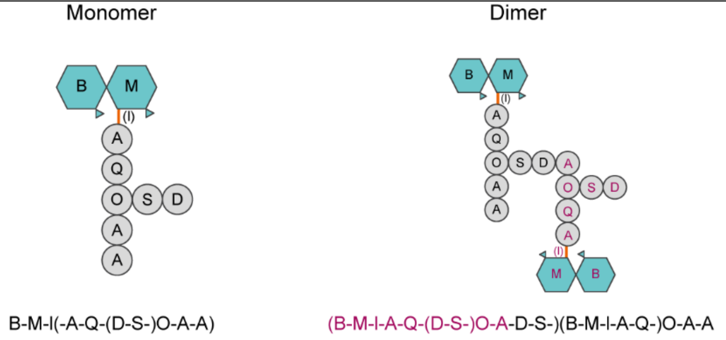


* The main products from PGN hydrolysis were labeled with bold font.

**Supplementary Table 7-1. Automated identification of *Bifidobacterium breve* ATCC 15700 muropeptides.**

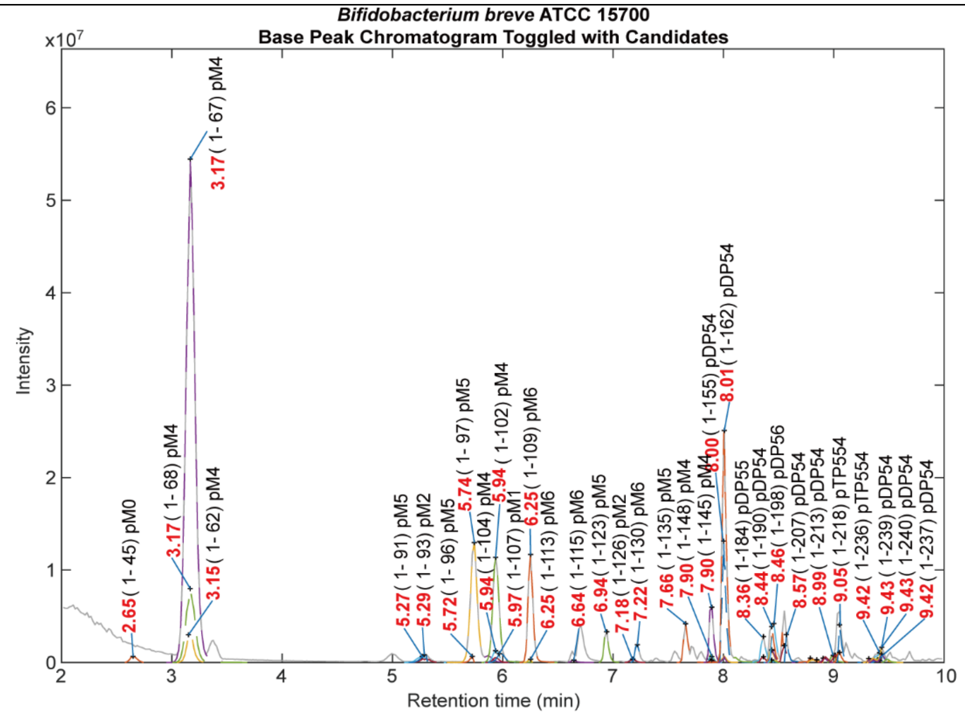


| Muropeptide structure (Main peaks) | RT (min) | Class | Observed MW (Da) | Theoretical MW (Da) | Δppm |
|---|---|---|---|---|---|
| **B-M-l(-A-Q-(G-)K)** | **3.17** | **Monomer** | **882.4182** | **882.4182** | **-0.03** |
| B-M-l(-A-Q-(G-)K-A) | 5.74 | Monomer | 953.4545 | 953.4553 | 0.90 |
| B-M-l(-A-E-(G-)K) | 5.94 | Monomer | 883.4031 | 883.4022 | -0.98 |
| B-M-l(-A-Q-(G-)K-A-A) | 6.25 | Monomer | 1024.4946 | 1024.4924 | -2.16 |
| **Tri-Tetra** | **8.01** | **Dimer** | **1817.8616** | **1817.8630** | **0.76** |
| Tetra-Tetra | 8.36 | Dimer | 1888.9031 | 1888.9001 | -1.60 |
| Tetra-Penta | 8.46 | Dimer | 1959.9376 | 1959.9372 | -0.20 |
| Tri-Tetra-Tetra | 9.05 | Trimer | 2753.3167 | 2753.3077 | -3.27 |
| Inferred types of peptide cross-links: 3-3 crosslink /4-3 crosslink | | | | | |

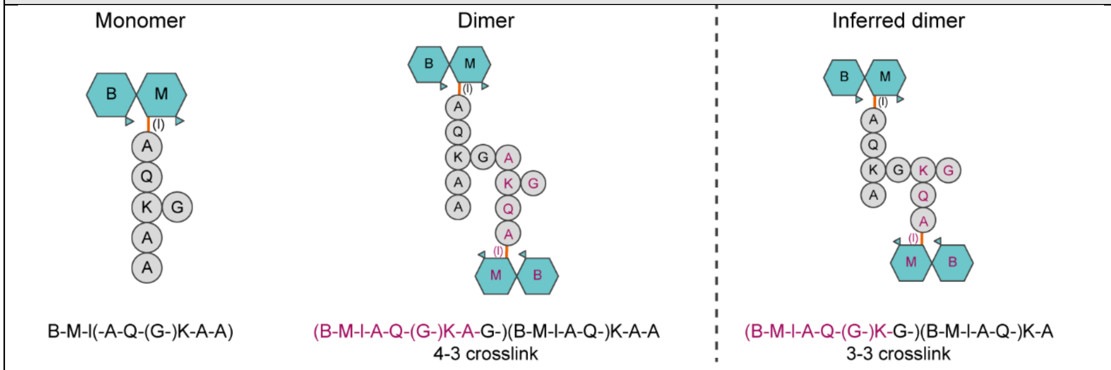


* The main products from PGN hydrolysis were labeled with bold font.

**Supplementary Table 7-2. Automated identification of *Bifidobacterium breve* CSCC 1900 muropeptides.**

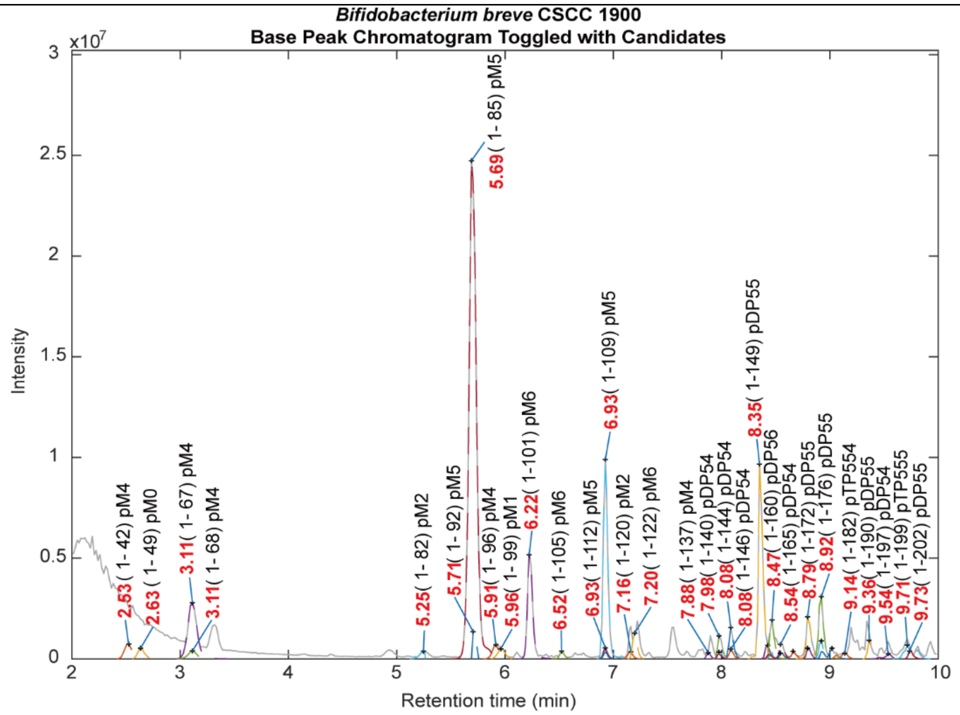


| Muropeptide structure (Main peaks) | RT (min) | Class | Observed MW (Da) | Theoretical MW (Da) | Δppm |
|---|---|---|---|---|---|
| B-M-l(-A-Q-(G-)K) | 3.11 | Monomer | 882.4184 | 882.4182 | -0.17 |
| **B-M-l(-A-Q-(G-)K-A)** | **5.69** | **Monomer** | **953.4542** | **953.4553** | **1.13** |
| B-M-l(-A-Q-(G-)K-A-A) | 6.22 | Monomer | 1024.4927 | 1024.4924 | -0.22 |
| **B-M-l(-A-E-(G-)K-A)** | **6.93** | **Monomer** | **954.4402** | **954.4393** | **-0.89** |
| Tri-Tetra | 8.08 | Dimer | 1817.8625 | 1817.8630 | 0.25 |
| **Tetra-Tetra** | **8.35** | **Dimer** | **1888.9019** | **1888.9001** | **-0.96** |
| Tetra-Penta | 8.47 | Dimer | 1959.9361 | 1959.9372 | 0.53 |
| Tetra-Tetra-Tetra (E, 2Q) | 9.71 | Trimer | 2825.3220 | 2825.3288 | 2.41 |

Inferred types of peptide cross-links: 4-3 crosslink /3-3 crosslink

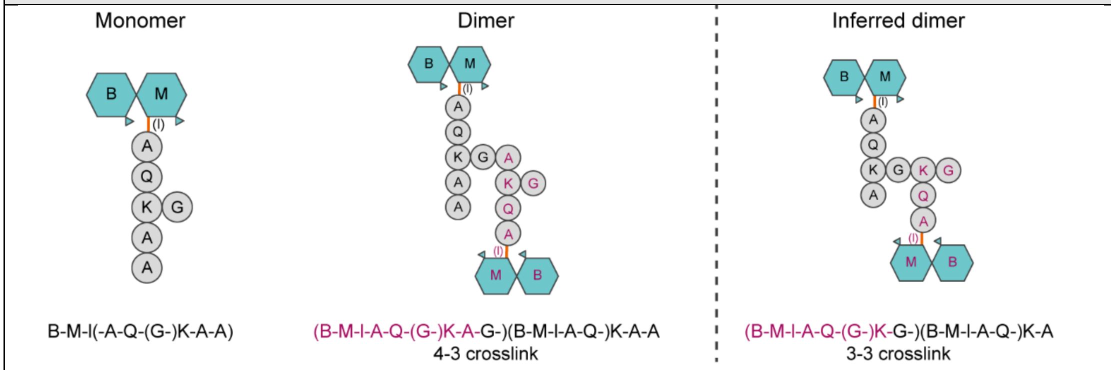


* The main products from PGN hydrolysis were labeled with bold font.

**Supplementary Table 7-3. Automated identification of *Bifidobacterium breve* ATCC 15698 muropeptides.**

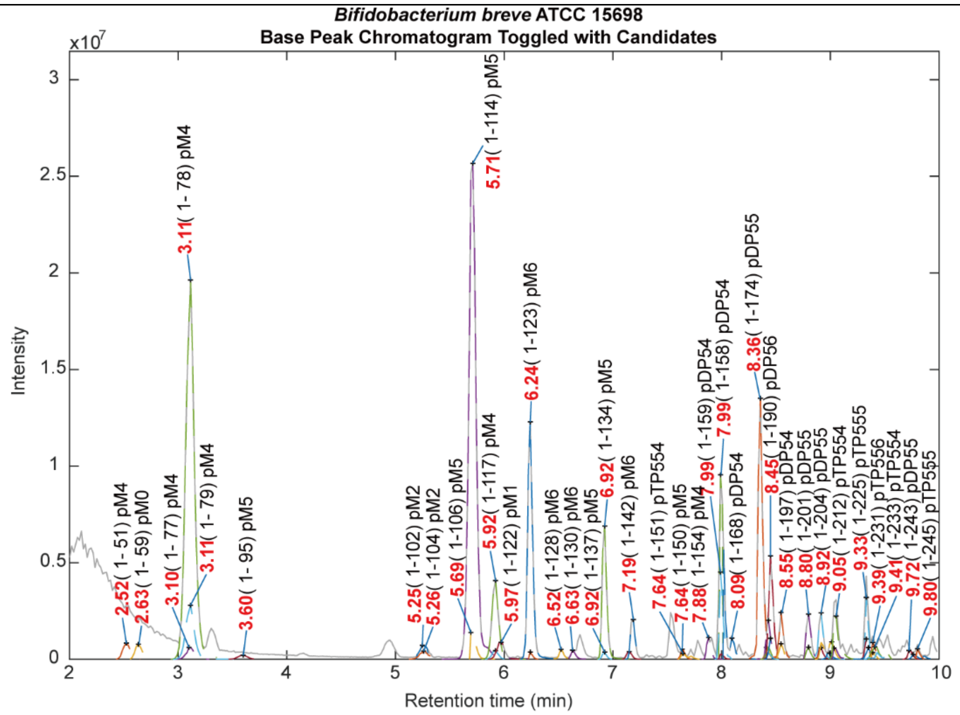


| Muropeptide structure (Main peaks) | RT (min) | Class | Observed MW (Da) | Theoretical MW (Da) | Δppm |
|---|---|---|---|---|---|
| **B-M-l(-A-Q-(G-)K)** | **3.11** | **Monomer** | **882.4185** | **882.4182** | **-0.30** |
| **B-M-l(-A-Q-(G-)K-A)** | **5.71** | **Monomer** | **953.4542** | **953.4553** | **1.17** |
| **B-M-l(-A-Q-(G-)K-A-A)** | **6.24** | **Monomer** | **1024.4943** | **1024.4924** | **-1.86** |
| B-M-l(-A-E-(G-)K-A) | 6.92 | Monomer | 954.4406 | 954.4393 | -1.28 |
| **Tri-Tetra** | **7.99** | **Dimer** | **1817.8574** | **1817.8630** | **3.04** |
| **Tetra-Tetra** | **8.36** | **Dimer** | **1888.9027** | **1888.9001** | **-1.37** |
| **Tetra-Penta** | **8.45** | **Dimer** | **1959.9366** | **1959.9372** | **0.30** |
| Tri-Tetra-Tetra | 9.05 | Trimer | 2753.3143 | 2753.3077 | -2.37 |

Inferred types of peptide cross-links: 4-3 crosslink /3-3 crosslink

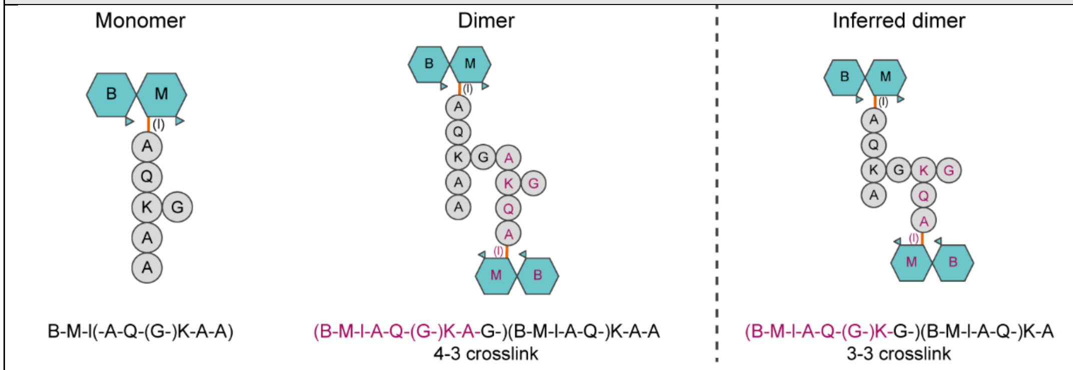


* The main products from PGN hydrolysis were labeled with bold font.

**Supplementary Table 8-1. Automated identification of *Bifidobacterium longum* ATCC 15707 muropeptides.**

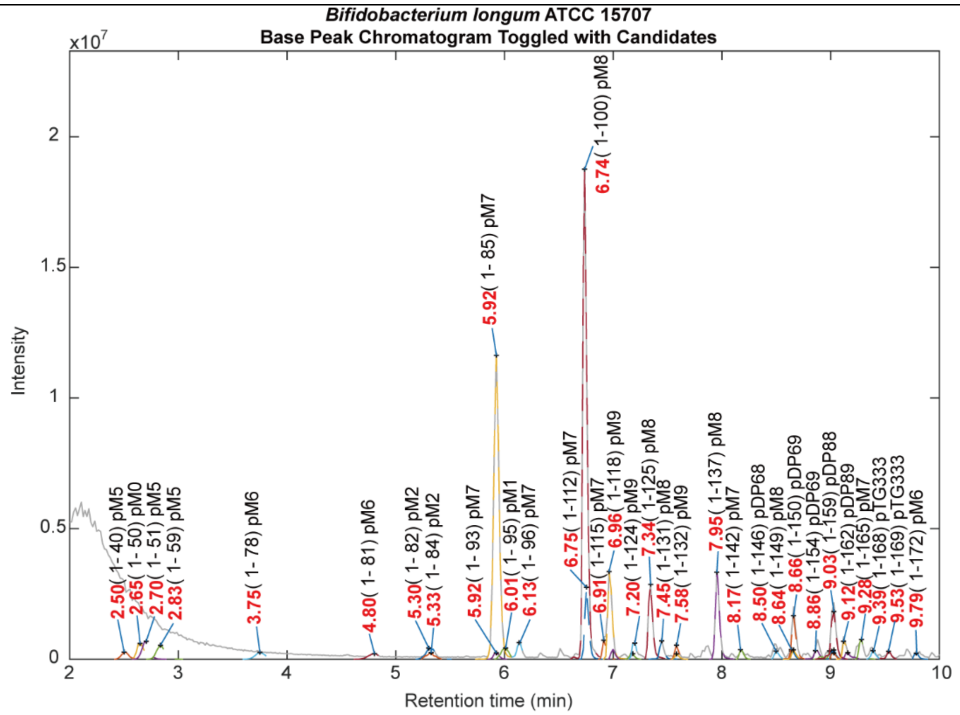


| Muropeptide structure (Main peaks) | RT (min) | Class | Observed MW (Da) | Theoretical MW (Da) | Δppm |
|---|---|---|---|---|---|
| **B-M-l(-A-Q-(A-T-A-S-)O)** | **5.92** | **Monomer** | **1141.5359** | **1141.5350** | **-0.73** |
| **B-M-l(-A-Q-(A-T-A-S-)O-A)** | **6.74** | **Monomer** | **1212.5727** | **1212.5721** | **-0.50** |
| B-M-l(-A-E-(A-T-A-S-)O) | 6.75 | Monomer | 1142.5162 | 1142.5190 | 2.50 |
| B-M-l(-A-Q-(A-T-A-S-)O-A-A) | 6.96 | Monomer | 1283.6086 | 1283.6093 | 0.48 |
| **B-M-l(-A-E-(A-T-A-S-)O-A)** | **7.34 /7.95** | **Monomer** | **1213.5562** | **1213.5562** | **-0.04** |
| Tri-Tetra | 8.66 | Dimer | 2336.1003 | 2336.0854 | -6.40 |
| **Tetra-Tetra** | **9.03** | **Dimer** | **2407.1369** | **2407.1337** | **-1.34** |
| Tetra-Penta | 9.12 | Dimer | 2478.1694 | 2478.1708 | 0.59 |
| Inferred types of peptide cross-links: 4-3 crosslink /3-3 crosslink | | | | | |

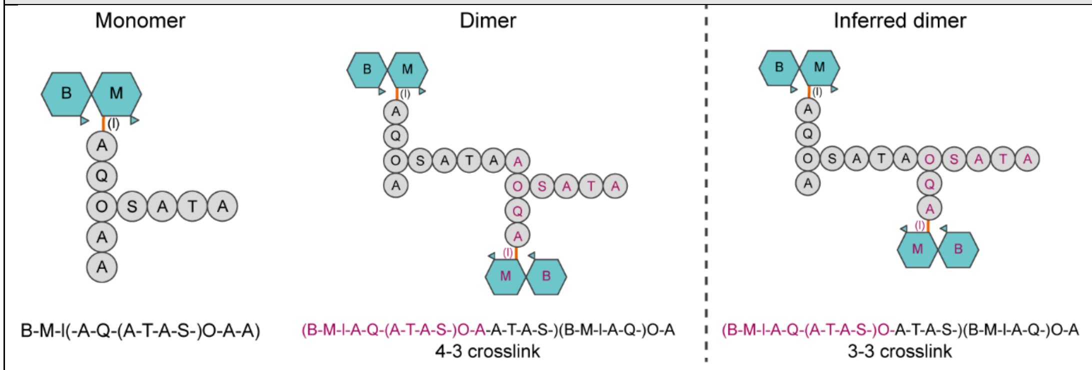


* The main products from PGN hydrolysis were labeled with bold font.

**Supplementary Table 8-2. Automated identification of *Bifidobacterium longum* CSCC 1901 muropeptides.**

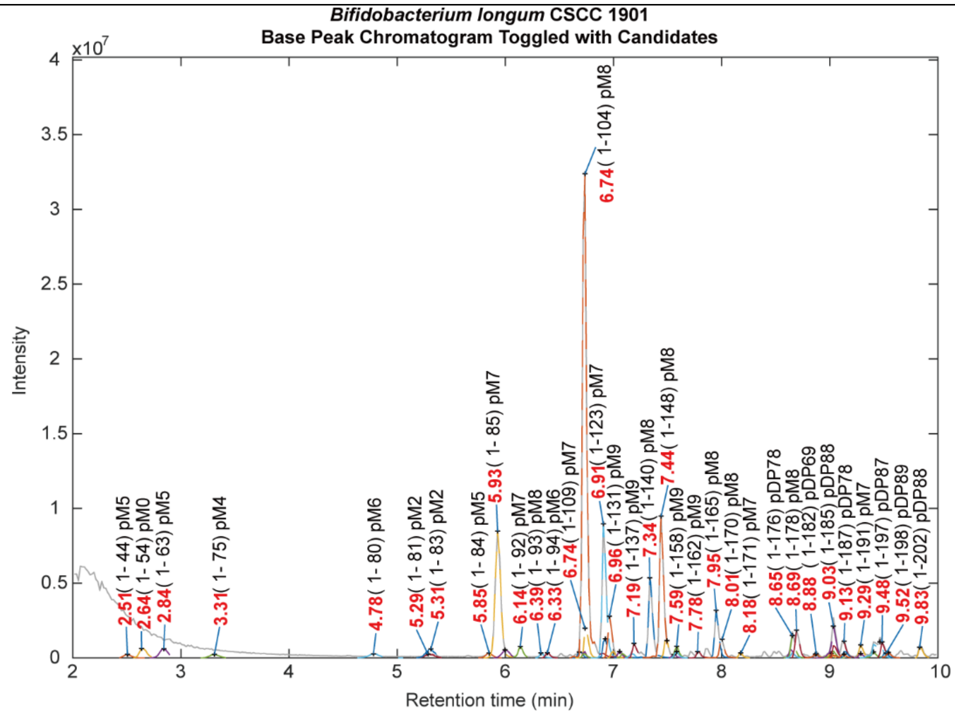


| Muropeptide structure (Main peaks) | RT (min) | Class | Observed MW (Da) | Theoretical MW (Da) | Δppm |
|---|---|---|---|---|---|
| **B-M-l(-A-Q-(A-T-A-S-)O)** | **5.93** | **Monomer** | **1141.5362** | **1141.5350** | **-0.99** |
| **B-M-l(-A-Q-(A-T-A-S-)O-A)** | **6.74** | **Monomer** | **1212.5743** | **1212.5721** | **-1.80** |
| B-M-l(-A-E-(A-T-A-S-)O) | 6.74 | Monomer | 1142.5159 | 1142.5190 | 2.72 |
| **B-M-l(-A-Q-(A-T-A-S-)K)** | **6.91** | **Monomer** | **1155.5484** | **1155.5507** | **1.96** |
| B-M-l(-A-Q-(A-T-A-S-)O-A-A) | 6.96 | Monomer | 1283.6095 | 1283.6093 | -0.18 |
| B-M-l(-A-E-(A-T-A-S-)O-A) | 7.34 | Monomer | 1213.5562 | 1213.5562 | -0.06 |
| **B-M-l(-A-Q-(A-T-A-S-)K-A)** | **7.44** | **Monomer** | **1226.5878** | **1226.5878** | **0.01** |
| Tetra-Tetra | 9.03 | Dimer | 2407.1380 | 2407.1337 | -1.76 |
| Inferred types of peptide cross-links: 4-3 crosslink /3-3 crosslink | | | | | |

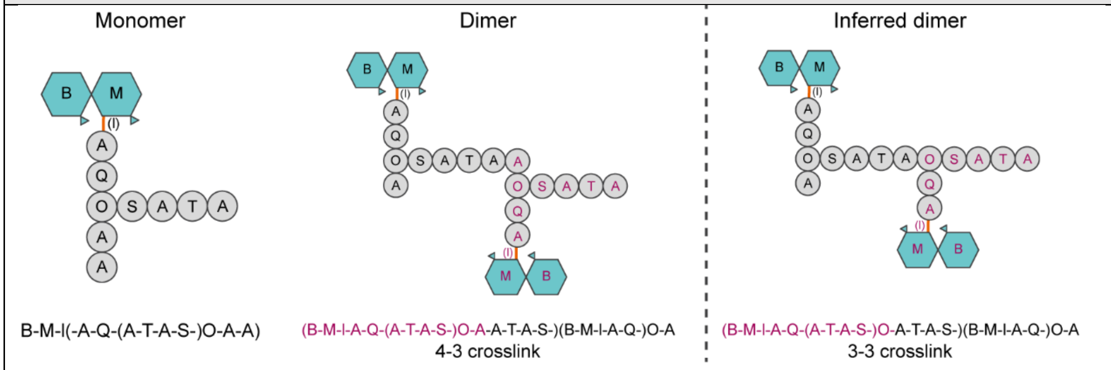


* The main products from PGN hydrolysis were labeled with bold font.

**Supplementary Table 8-3. Automated identification of *Bifidobacterium longum* ATCC 15697 muropeptides.**

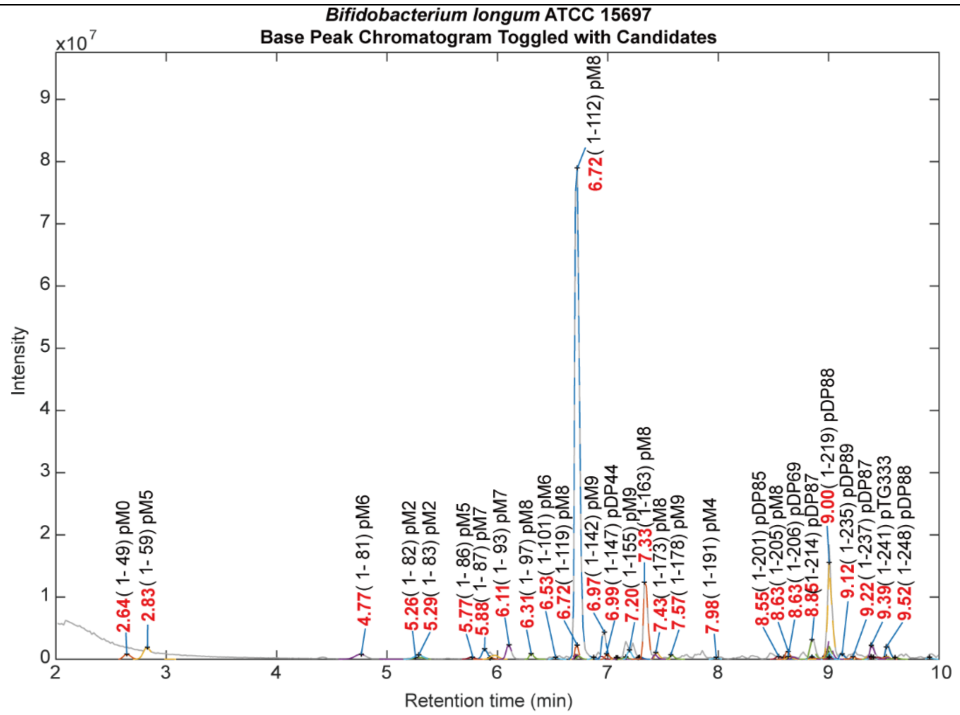


| Muropeptide structure (Main peaks) | RT (min) | Class | Observed MW (Da) | Theoretical MW (Da) | Δppm |
|---|---|---|---|---|---|
| B-M-l(-A-Q-(S-)O-A) | 2.83 | Monomer | 969.4481 | 969.4502 | 2.25 |
| **B-M-l(-A-Q-(A-T-A-S-)O-A)** | **6.72** | **Monomer** | **1212.5748** | **1212.5721** | **-2.21** |
| B-M-l(-A-Q-(A-T-A-S-)O-A-A) | 6.97 | Monomer | 1283.6092 | 1283.6093 | 0.07 |
| **B-M-l(-A-E-(A-T-A-S-)O-A)** | **7.33** | **Monomer** | **1213.5605** | **1213.5562** | **-3.60** |
| Tri-Tetra | 8.85 | Dimer | 2336.1073 | 2336.0966 | -4.59 |
| **Tetra-Tetra** | **9.00** | **Dimer** | **2407.1438** | **2407.1337** | **-4.19** |
| Inferred types of peptide cross-links: 4-3 crosslink /3-3 crosslink | | | | | |

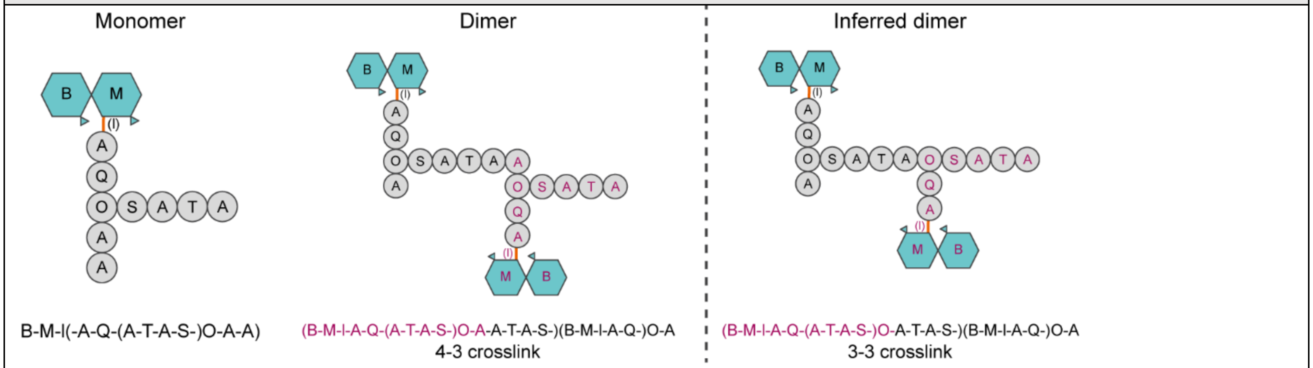


* The main products from PGN hydrolysis were labeled with bold font.

**Supplementary Table 9. Automated identification of *Bacteroides fragilis* ATCC 25285 muropeptides.**

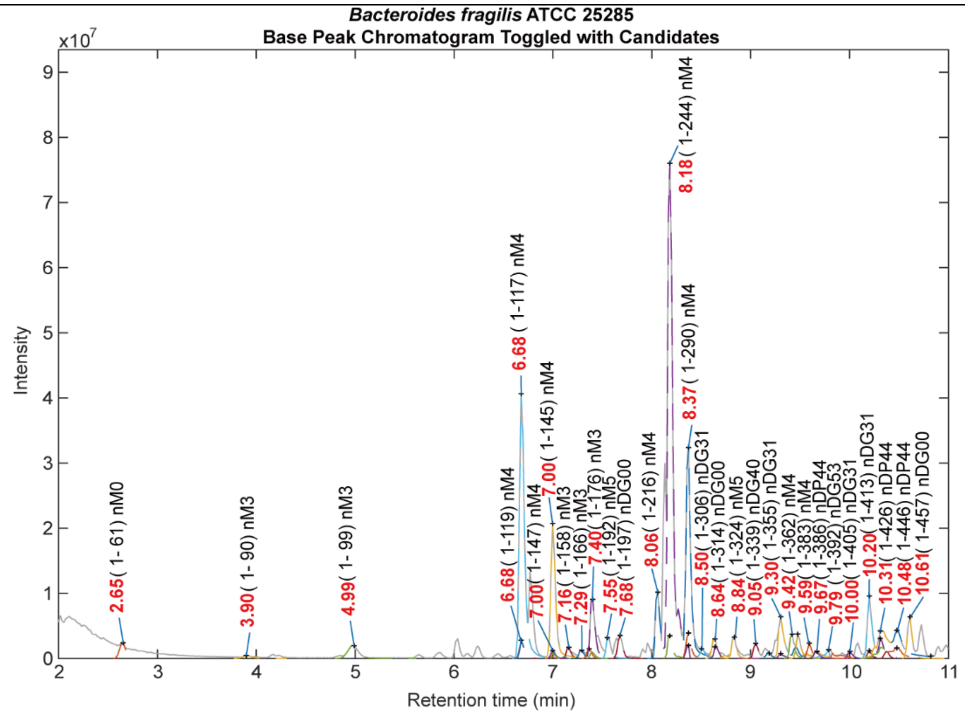


| Muropeptide structure (Main peaks)<br>Red: glycosidic bond | RT (min) | Class | Observed MW (Da) | Theoretical MW (Da) | Δppm |
|---|---|---|---|---|---|
| B-M-l(-A-E-H) | 4.99 | Monomer | 870.3708 | 870.3706 | -0.26 |
| **B-M-l(-A-E-H-A)** | **6.68/ 7.00** | **Monomer** | **941.4093** | **941.4077** | **-1.71** |
| B-Z-l(-A-E-H) | 7.40 | Monomer | 912.3811 | 912.3812 | 0.01 |
| **B-Z-l(-A-E-H-A)** | **8.18/ 8.37** | **Monomer** | **983.4170** | **983.4183** | **1.25** |
| Tri-Mono (m, Z) | 9.30 | Dimer | 1461.5970 | 1461.5981 | 0.75 |
| **Tri-Mono (z, Z)** | **10.20** | **Dimer** | **1503.6092** | **1503.6087** | **-0.33** |
| Tetra-Tetra (M, Z) | 10.48 | Dimer | 1906.8151 | 1906.8154 | 0.16 |
| Inferred types of peptide cross-links: 4-3 crosslink | | | | | |

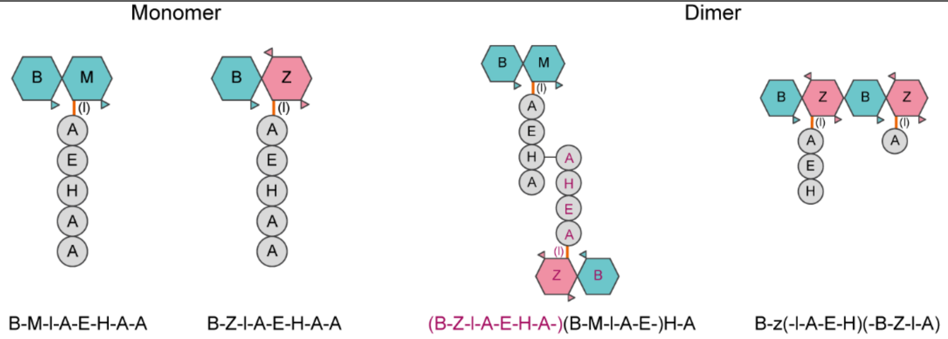


* The main products from PGN hydrolysis were labeled with bold font.

**Supplementary Table 10. Automated identification of *Bacteroides ovatus* ATCC 8483 muropeptides.**

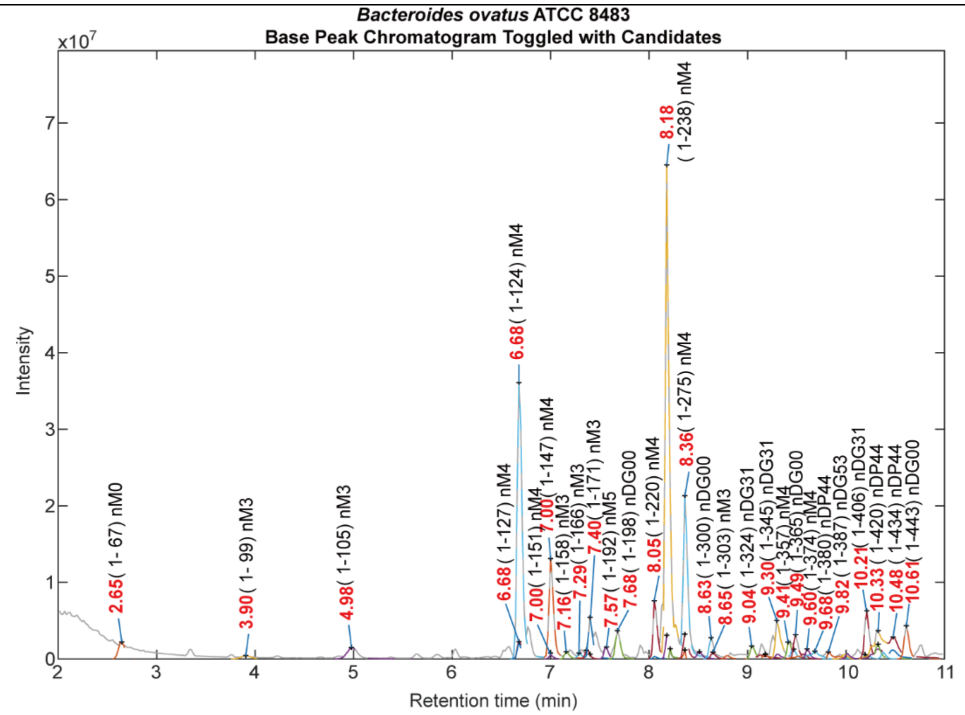


| Muropeptide structure (Main peaks)<br>Red: glycosidic bond | RT (min) | Class | Observed MW (Da) | Theoretical MW (Da) | Δppm |
|---|---|---|---|---|---|
| B-M-l(-A-E-H) | 4.98 | Monomer | 870.3700 | 870.3706 | 0.67 |
| **B-M-l(-A-E-H-A)** | **6.68/ 7.00** | **Monomer** | **941.4078** | **941.4077** | **-0.14** |
| B-Z-l(-A-E-H) | 7.40 | Monomer | 912.3808 | 912.3812 | 0.40 |
| **B-Z-l(-A-E-H-A)** | **8.18/ 8.36** | **Monomer** | **983.4169** | **983.4183** | **1.34** |
| Tri-Mono (m, M) | 8.36 | Dimer | 1419.5836 | 1419.5876 | 2.81 |
| Tri-Mono (m, Z) | 9.30 | Dimer | 1461.5969 | 1461.5981 | 0.85 |
| **Tri-Mono (z, Z)** | **10.21** | **Dimer** | **1503.6084** | **1503.6087** | **0.18** |
| Tetra-Tetra (M, Z) | 10.48 | Dimer | 1906.8200 | 1906.8154 | -2.39 |
| Inferred types of peptide cross-links: 4-3 crosslink | | | | | |

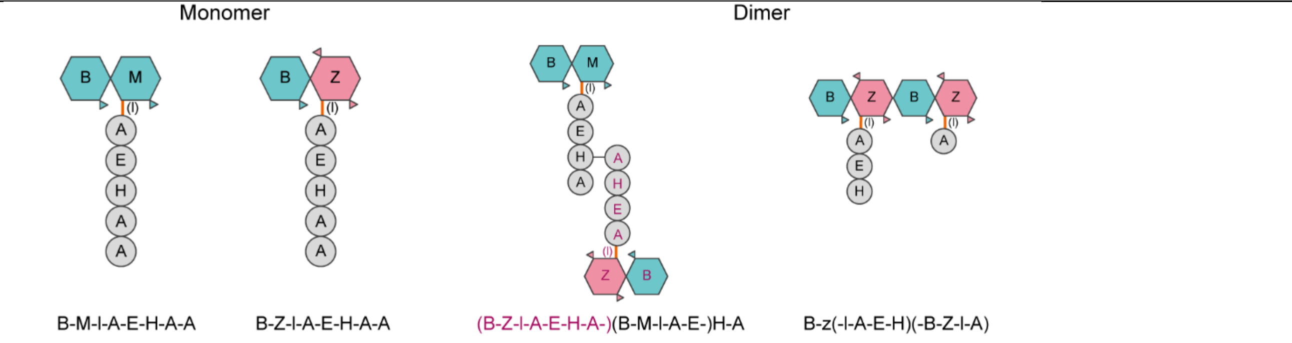


* The main products from PGN hydrolysis were labeled with bold font.

**Supplementary Table 11. Automated identification of *Bacteroides thetaiotaomicron* ATCC 29741 muropeptides.**

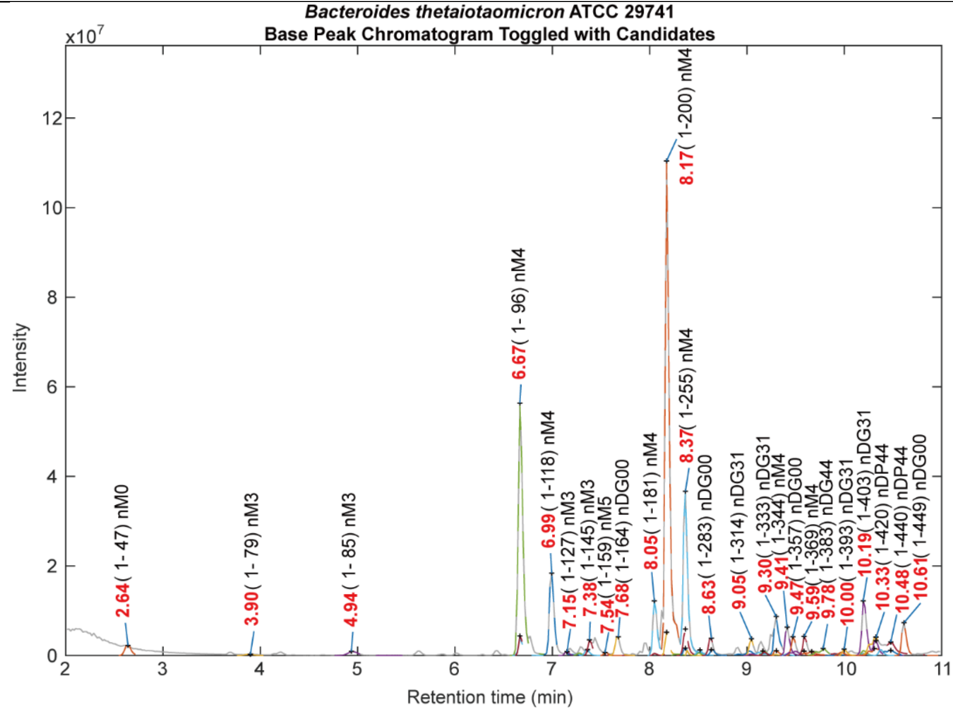


| Muropeptide structure (Main peaks)<br>Red: glycosidic bond | RT (min) | Class | Observed MW (Da) | Theoretical MW (Da) | Δppm |
|---|---|---|---|---|---|
| **B-M-l(-A-E-H-A)** | **6.67/ 6.99** | **Monomer** | **941.4096** | **941.4077** | **-2.04** |
| **B-Z-l(-A-E-H-A)** | **8.17/ 8.37** | **Monomer** | **983.4171** | **983.4183** | **1.16** |
| Tri-Mono (m, M) | 8.37 | Dimer | 1419.5847 | 1419.5876 | 2.04 |
| Tri-Mono (m, Z) | 9.30 | Dimer | 1461.5982 | 1461.5981 | -0.02 |
| B-U-l(-A-E-H-A) | 9.41 | Monomer | 921.3828 | 921.3815 | -1.42 |
| **Tri-Mono (z, Z)** | **10.19** | **Dimer** | **1503.6091** | **1503.6087** | **-0.23** |
| Tetra-Tetra (M, Z) | 10.33 | Dimer | 1906.8205 | 1906.8154 | -2.67 |
| B-z(-l)(-B-Z-l), glycan chain | 10.61 | Dimer | 1060.4072 | 1060.4071 | -0.09 |
| Inferred types of peptide cross-links: 4-3 crosslink | | | | | |

Schematic representation

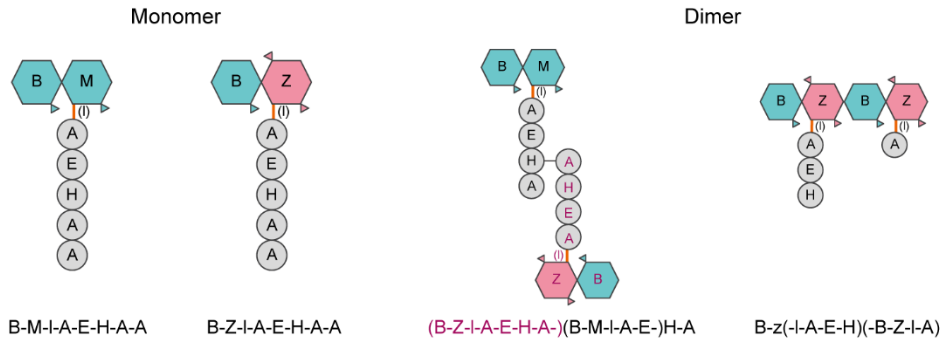


* The main products from PGN hydrolysis were labeled with bold font.

**Supplementary Table 12. Automated identification of *Akkermansia muciniphila* ATCC BAA-835 muropeptides.**

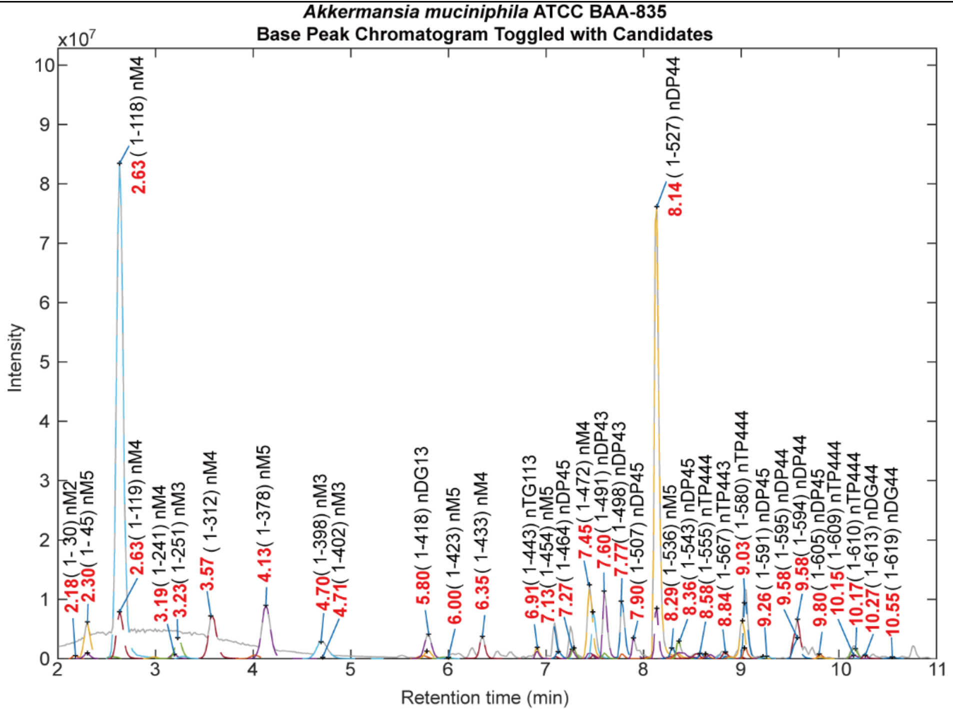


| Muropeptide structure (Main peaks) | RT (min) | Class | Observed MW (Da) | Theoretical MW (Da) | Δppm |
|---|---|---|---|---|---|
| **C-M-l(-A-E-H-A)** | **2.63/ 3.57** | **Monomer** | **899.39560** | **899.3971** | **1.71** |
| C-M-l(-A-E-H-A-A) | 4.13 | Monomer | 970.43364 | 970.4343 | 0.63 |
| B-M-l(-A-E-H-A) | 7.45 | Monomer | 941.40724 | 941.4077 | 0.50 |
| Tri-Tetra | 7.60/ 7.77 | Dimer | 1709.74873 | 1709.7466 | -1.25 |
| **Tetra-Tetra** | **8.14** | **Dimer** | **1780.78675** | **1780.7837** | **-1.71** |
| Tetra-Tetra-Tetra | 9.03 | Dimer | 2662.17591 | 2662.1703 | -2.11 |
| Tetra-Tetra (B, C) | 9.58 | Dimer | 1822.79354 | 1822.7943 | 0.41 |

Inferred types of peptide cross-links: 4-3 crosslink/ 3-3 crosslink

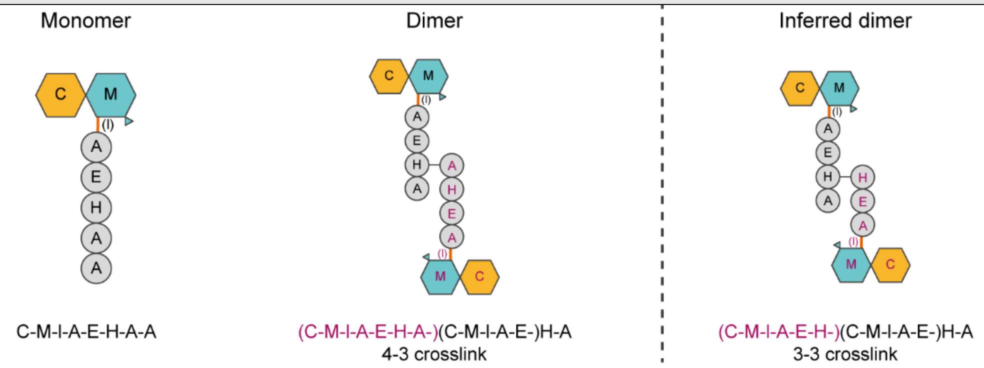


* The main products from PGN hydrolysis were labeled with bold font.
